# Extended categorization of conjunction object stimuli decreases the latency of attentional feature selection and recruits orthography-linked ERPs

**DOI:** 10.1101/605360

**Authors:** Jonathan R. Folstein, Shamsi S. Monfared

## Abstract

The role of attention in driving perceptual expertise effects is controversial. The current study addressed the effect of training on ERP components related to and independent of attentional feature selection. Participants learned to categorize cartoon animals over six training sessions (8,800 trials) after which ERPs were recorded during a target detection task performed on trained and untrained stimulus sets. The onset of the selection negativity, an ERP component indexing attentional modulation, was about 60 ms earlier for trained than untrained stimuli. Trained stimuli also elicited centro-parietal N200 and N320 components that were insensitive to attentional feature selection. The scalp distribution and timecourse of these components were better matched by studies of orthography than object expertise. Source localization using eLORETA suggested that the strongest neural sources of the selection negativity were in right ventral temporal cortex whereas the strongest sources of the N200/N320 components were in left ventral temporal cortex, again consistent with the hypothesis that training recruited orthography related areas. Overall, training altered neural processes related to attentional selection, but also affected neural processes that were independent of feature selection.

## Introduction

Studies using several dependent measures have demonstrated that neural representations of objects of expertise, including cars (Gauthier, Skudlarski, Gore, & Anderson, 2000; Scott, Tanaka, Sheinberg, & Curran, 2008), birds (Gauthier et al., 2000; Scott, Tanaka, Sheinberg, & Curran, 2006), chessboards (Bilalić, Langner, Ulrich, & Grodd, 2011), x-rays (Bilalić, Grottenthaler, Nägele, & Lindig, 2014), novel objects (Gauthier, Tarr, Anderson, Skudlarski, & Gore, 1999; Jones et al., 2018), and words (Mei et al., 2013; Xue, Chen, Jin, & Dong, 2006), are different from other object categories in expert populations, suggesting that experience is an important variable driving those representations.

The role of attention in perceptual expertise is an important focus of research and debate. Some have argued that attention is a confounding factor in studies claiming that expertise recruits specialized modules like the FFA (Harel, Gilaie-Dotan, Malach, & Bentin, 2010; McKone, Kanwisher, & Duchaine, 2007). Others have proposed learned attention as a mechanism for holistic perception in perceptual expertise (Chua, Richler, & Gauthier, 2015). In general, learning to categorize objects is known to cause strategic changes in allocation of attention to category-relevant features and dimensions (Blair, Watson, Walshe, & Maj, 2009; Li, Ostwald, Giese, & Kourtzi, 2007). Furthermore, Jolicoeur, Gluck, and Kosslyn (1984) showed that subordinate level categorization, a hallmark of perceptual expertise, requires more detailed perceptual analysis than basic level categorization. Thus, the finding that expertise facilitates subordinate level categorization (Tanaka & Taylor, 1991) could imply facilitated ability to apply attention to objects of expertise.

Event related potentials (ERPs) offer another piece of evidence that perceptual expertise could result in altered attentional processes. In particular, the N250 is an expertise-linked component that has a postero-lateral scalp distribution and post-200ms onset consistent with an effect of top-down feedback to visual cortex (e.g. Ganis & Kosslyn, 2007). The timecourse and scalp distribution of the N250 is similar to the selection negativity, a component linked to attention (Daffner et al., 2012; Gledhill, Grimsen, Fahle, & Wegener, 2015; Hillyard & Anllo-Vento, 1998), which is known to be an important source of feedback modulating visual areas (e.g. Baldauf & Desimone, 2014; Gregoriou, Gotts, Zhou, & Desimone, 2009). In training studies, the N250 is enhanced by trained stimulus categories and consistently selective for stimuli that have been categorized at the subordinate level (blue jay vs. scrub jay) (Jones et al., 2018; Scott et al., 2006, 2008). The N250 is also larger to highly familiar, easily identified objects, such as non-target faces that have recently been targets (Gordon & Tanaka, 2011), one’s own face, own dog, or own car, than less familiar objects of the same categories (Pierce et al., 2011), suggesting that it is driven by recognition of objects and subordinate categories at the individual level. We argued previously (Folstein, Monfared, & Maravel, 2017) that N250 effects have plausible attention related explanations even in cases where they are not elicited by explicit target recognition. The training studies, for instance, elicit the N250 in a match-to-category task in which participants must evaluate subordinate level category membership (Scott et al., 2006, 2008). Experts could apply attention differently in such a task because they know which features are diagnostic for particular categories and, unlike novices, are likely to attentionally select them during the task with high precision.

Importantly, attention is not the only source of feedback to visual cortex. Price and Devlin (2011) proposed that activation in the VWFA was driven by automatically accessed **non-strategic feedback** from higher areas, facilitating recognition of the perceptual features of words. Consistent with this proposal words also sometimes elicit later vision-related ERP components, including an N250 (Carreiras, Duñabeitia, & Molinaro, 2009; Grainger, Kiyonaga, & Holcomb, 2006) and N200 (Ruz & Nobre, 2008; Zhang et al., 2012; Zhou, Yin, Zhang, & Zhang, 2016), both of which are associated with orthographic processing. These later components often have a more dorsal scalp distribution than the postero-lateral N250 effects observed in studies of perceptual expertise. Others have proposed orbitofrontal cortex and medial temporal cortex as other sources of rapidly accessed non-strategic feedback subserving visual object recognition (see also Bar, 2003; Barense, Ngo, Hung, & Peterson, 2012).

A previous study in our lab (Folstein, Monfared, et al., 2017) supported the possibility that the non-strategic feedback could contribute to the N250. A two-day category learning task resulted in an N250 learning effect that was additive with a selection negativity effect driven by attention to features shared with a target. Both components had postero-lateral scalp distributions and similar time courses, the selection negativity (SN) being significant from 190 to 380 milliseconds and the N250 from 270 to 300 ms. Because the components were additive and did not interact, we concluded that that neural effects related to perceptual training could be dissociated from at least some aspects of attentional selection. Furthermore, no effect training was observed on selection negativity, suggesting that training had little effect on attention. One limitation of the study was that participants were only trained for two sessions for far fewer trials than are usually employed in studies of perceptual expertise. A longer training period could be a more powerful manipulation, altering the effectiveness with which strategic attention is applied. As we will see, the longer training period had a major impact on late non-strategic components as well.

## The current study

The primary goal of the current study was to investigate whether a more extended period category learning, known to cause perceptual expertise (Gauthier & Tarr, 1997; Wong, Palmeri, & Gauthier, 2009), a) effects neural correlates of attentional selection and b) affects neural representations in a way that could be dissociated from effects of attention. Participants learned to categorize a set of cartoon animals while a second set was untrained. After training, ERPs were recorded while participants detected a single target exemplar from the trained or untrained set. Non-target exemplars shared between zero and four features with the target.

The selection negativity served as our operational definition of attentional feature selection. Each target feature contained in a given non-target stimulus was predicted to elicit a selection negativity component. Rather than attempting to isolate the SN component for each feature, we measured the cumulative selection negativity, which increases in amplitude with increasing numbers of target features. We have generally observed this increase to be linear (Folstein, Fuller, Howard, & DePatie, 2017; Folstein, Monfared, et al., 2017). As in our previous study, we predicted that, if training alters attentional feature selection, the effect of training should interact with the selection negativity, altering the slope of the relationship between the SN and number of target features (Folstein, Fuller, et al., 2017). The ways that training induced changes to attentional selection might affect the slope of the SN are reviewed in detail in our previous paper (Folstein, Monfared, et al., 2017); in general more effective feature selection is predicted to increase the slope, whereas a transition from selecting single features to conjunctions of features is predicted to make the slope less linear. Smid, Jakob, and Heinze (1999) observed that salient, easy to detect visual features elicited earlier, more long lasting selection negativity components. Thus, if training has the effect of making features easier to discriminate, then training should result in an earlier SN onset.

Several aspects of our stimulus set suggested that it should be effective in eliciting the standard N170 and N250 expertise effects. First, the stimuli were animals, which are known to be strongly represented in the FFA (e.g. McGugin, Gatenby, Gore, & Gauthier, 2012). Second, the categorization task required “fine grained” discriminations such that stimuli that shared many features had to be assigned to different categories. Third, our previous study (Folstein, Monfared, et al., 2017), which used only two days of training, was successful in eliciting a postero-lateral N250 training effect.

Two aspects of our stimuli are unusual for object training studies. First, the animals are constructed from conjunctions of interchangable parts (e.g. head, body, legs, tail), each with two binary values. The purpose of this characteristic is to make parametric manipulation of similarity convenient in a feature theoretic stimulus set, a strategy that has been used in many previous studies of categorization (e.g. Allen & Brooks, 1991; Medin & Schaffer, 1978). However, interchangeable parts are unrealistic in the sense that most real subordinate level object categories do not share exact duplicates of features – rather they have **similar** features. For instance, Blue jays, Stellar Jays, and Cardinals, all have crests but the shape of crest differs somewhat between species. A closely related point is that objects are also not defined purely by conjunctions of features, but instead have **individuating** features with various degrees of salience, such as the distinctive blue, black and white pattern on a Blue Jay’s back. A second potentially important characteristic of our stimulus set is that it is 2D rather than shape-from-shading and the stimuli are defined by black outlines (Figure 1).

**Figure 1.**
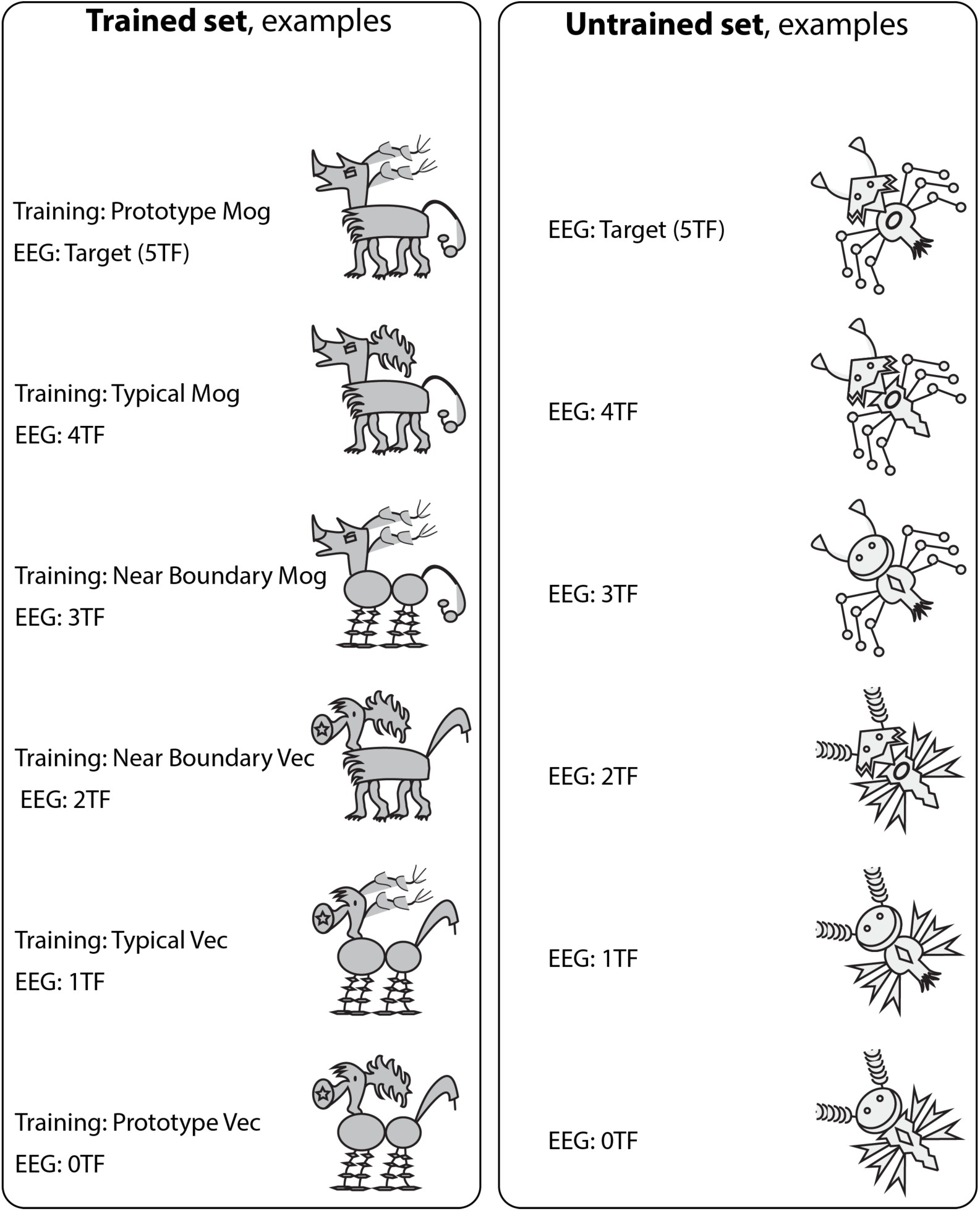
Examples from each the two stimulus sets. Which set was trained was counterbalanced across participants.

These characteristics make the stimuli we used similar to words. Words are defined by conjunctions of interchangeable features (letters) and contain line junctions, which were hypothesized by Dehaene (e.g. Dehaene & Cohen, 2011) to recruit the visual word form area (VWFA). We call attention to this characteristic here to prepare the reader for an unexpected finding. Whereas we again observed dissociable effects of training and attention, the scalp distribution of the training effect changed and was now highly inconsistent with N250 training effects observed in past studies of perceptual expertise for objects (Jones et al., 2018; Scott et al., 2006, 2008). Instead, the strongly centro-parietal component we observed was more consistent with certain studies of orthography, which we review below. This is particularly interesting because the stimuli we used were animals and nothing about the training we used invited participants to think of them as orthographic stimuli.

We also explored the effect of fast and slow stimulus onset asynchronies (SOA) on effects of training and attention. The main reason for our manipulation of SOA was to prepare for a later fMRI experiment, which requires a slower SOA (two seconds or more) than used in our previous experiment. Overall, SOA did affect some ERPs of interest, such as the N170, but did not interact with effects of training or attention.

## Methods

### Subjects

Twenty seven students from Florida State University below the age of 30 were recruited for the experiment. Two were excluded because they did not pass the screening test; 6 were discarded because they failed to complete all training sessions, and 1 was dropped due to a technical error during training, leaving 16 participants that completed the entire experiment, eight of which were female. Participants reported vision corrected to 20/20 and no history of head injury or psychiatric illness.

### Stimuli

Stimuli were composed of two sets of cartoon aliens: alien horses and alien bugs (Figure 1), spanning approximately 3.5 degrees of visual angle. For each participant, one set was trained and the other set was untrained, counterbalanced across participants. The 32 exemplars in each set (Figure S1) were constructed by factorially combining five two-valued features (horses: head, main, body, legs, and tail; bugs: antennae, head, body, legs, markings). Each set could be divided into two categories using a “3-out-of-5 rule”, according to which an exemplar was in a category if three out of the five features were associated with that category. Each category included one prototype with all five features, five “typical” members with four out of five features, and ten “near boundary” members with three out of five features. The feature values associated with each category were randomly assigned for each participant with the exception that each category-feature assignment was used as “trained” for one participant and “untrained” for a second participant.

## Procedure

The procedures prior to the EEG session were identical to that used by (Folstein, Monfared, et al., 2017) except that six training sessions were conducted rather than two (Figure 2).

**Figure 2.**
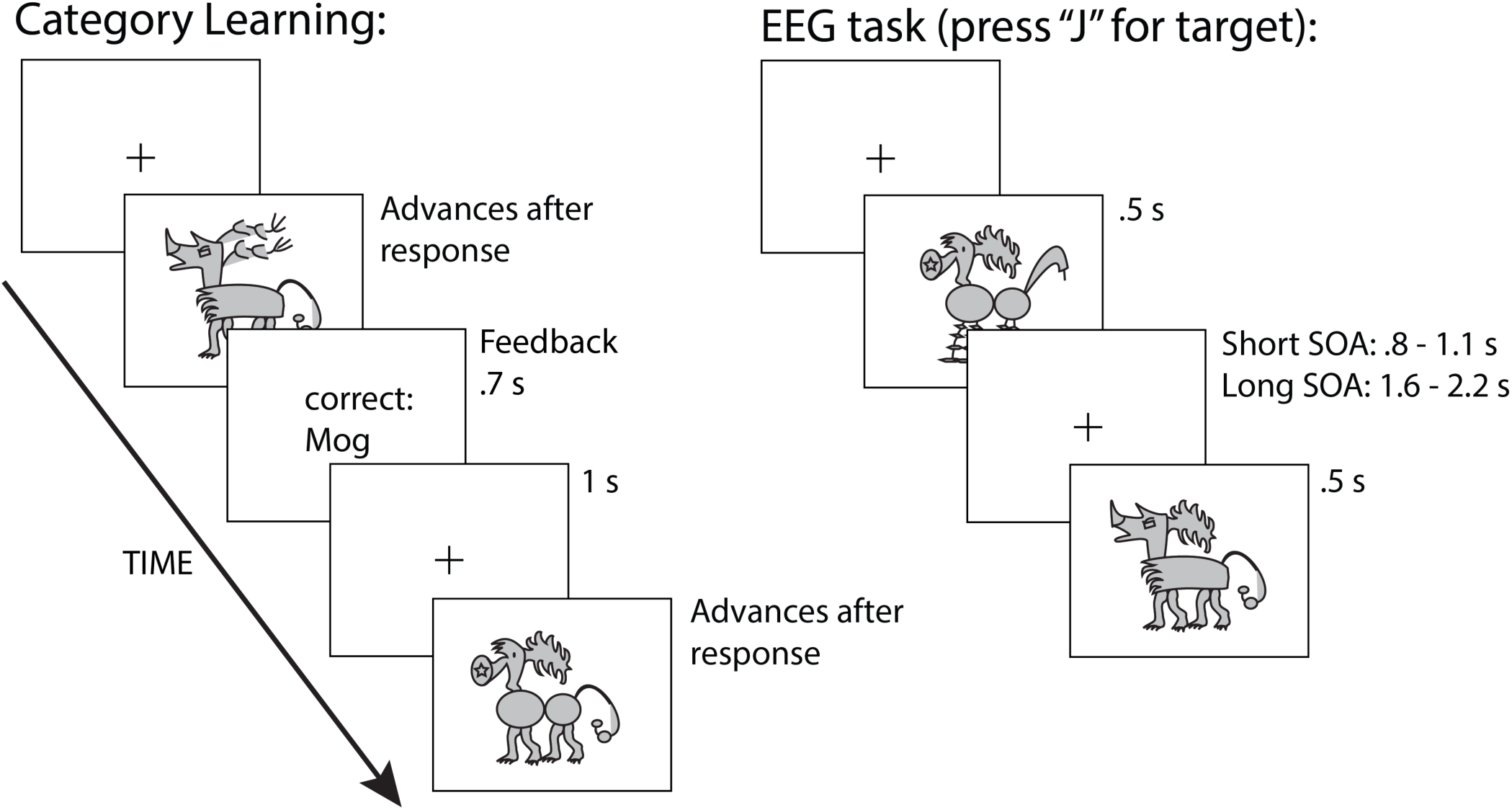
Category learning task and target detection task performed during EEG recording.

### Screening task

Prior to the first training session, participants performed four blocks of the target detection task that included both stimulus sets. Two participants failed to pass the screening with less than 75% correct responses, but after this we decided to allow all participants to advance to training even if they failed so that participants could be run more rapidly. As a result, one participant that failed the screening was included in the experiment. All participants received the screening to maintain equal exposure to the stimuli.

### Category learning

The category learning task included 1) a study phase during which participants learned the categorization rule for the trained set and advanced through labeled exemplars at their own pace and 2) a training phase during which they categorized unlabeled exemplars and received feedback on their accuracy. Participants were allowed to return to the study phase during training if they were not confident that they understood the categorization rule.

The study phase was identical to Folstein, Monfared, et al. (2017). Briefly, participants 1) reviewed the stimulus features 2) received verbal instructions describing the 2-out-of-3 rule 3) viewed the prototypes 4) viewed the exemplars at their own pace as many times as they wished; at this stage the stimuli appeared along with category labels (e.g. Mog) and feature labels (e.g. Mog head) 5) categorized unlabeled exemplars in random order with feedback (Figure 2); this stage included 8,800 trials (session 1: 1120 trials; sessions 2 to 6: 1536 trials each). See supplemental materials for a more detailed description.

### Target detection task

During EEG recording, participants responded to rare target stimulus with a button press, ignoring all other stimuli. Trained stimuli were presented during four runs (360 trials per run) and untrained stimuli were presented during four runs, with stimulus set alternating every two runs, order counterbalanced across subjects. Half of the runs in each training condition used a fast stimulus onset asynchrony (SOA) and half used a slow SOA, alternating every run with order counterbalanced across conditions. The target was always the Mog prototype (recall that the particular stimuli assigned to be the Mog and Vec prototypes were randomized across pairs of participants). Non-target stimuli shared 4 (typical Mog), 3 (near-boundary Mog), 2 (near-boundary Vec), 1 (Typical Vec), or 0 (Vec prototype) features with the target. Each non-prototype condition included 5 exemplars, pseudo-randomly chosen to preserve equal frequency for all features. Prototypes (targets) and Vec prototypes each had one exemplar, presented five times per block. Target probability of 16.67%.

For the fast SOA condition interstimulus interval was jittered randomly between 800, 900, 1000, and 1100 ms; ISIs for the slow SOA condition were 1600, 1800, 2000, and 2200 ms. Participants received a break every 60 trials, at which point they received a reminder of the target stimulus and instructions to keep their eyes on the location of the fixation cross. Responses between 350 and 2000 ms after the target were counted as correct.

## EEG data acquisition

EEG was recorded using a 64 channel BrainAmpDC amplifier (Brain Products), running in “AC mode” and gel cap with 62 equidistant AgCl electrodes referenced to the right mastoid. Two additional electrodes were placed under the eyes, 15% of interaural distance lateral and 20% of inion-nasion distance down from FPz. Electrode impedance was reduced to less than 5 kohms prior to recording. Participants were seated in an electrically shielded, sound attenuated chamber 4 feet from the monitor. Stimuli were presented on a 23.6 inch 120-Hz ViewPixx white LED backlight monitor with rise and fall times comparable to CRT monitors. EEG was monitored during recording and electrodes adjusted as needed throughout the session.

## EEG data analysis

All data analysis was conducted in Matlab using EEGLAB (Delorme & Makeig, 2004) and ERPLAB (Lopez-Calderon & Luck, 2014) toolboxes. Artifact rejection was conducted in three stages. First, large artifacts were removed automatically using the EEGLAB function clean_rawdata, which uses artifact subspace reconstruction to detect artifacts. All parameters except time window removal were set to −1 so that EEG was removed but not corrected; the window removal parameter was set to between .3 and .4. The results of this step were evaluated by eye to ensure that only large artifacts were removed. Second, Independent Components Analysis (ICA) was run and ICs corresponding to blinks were removed. After data epoching, a second round of artifact rejection was conducted using modified versions of the, pop_artmwppth, pop_artdiff, and pop_rejtrend tests available in ERPLAB and EEGLAB, which test specific channels for large amplitude shifts within a trial, amplitude shifts between temporally adjacent samples, and gradual linear trends, respectively. The results of these tests were evaluated by eye for each participant so that their parameters could be adjusted to optimally reject artifacts and accept good trials. After all artifact stages and removing inaccurate trials, an average of between 74.7 and 88.6 trials remained in each cell of the total 5 × 2 × 2 (0 to 5 target features; trained vs. untrained; long SOA vs. short SOA).

Our choice of reference was influenced by our desire to make our results comparable to other studies of the N250, particularly those related to the effect of training and expertise (Jones et al., 2018; Pierce et al., 2011; Scott et al., 2006, 2008). These studies all used average reference and high density caps. Liu et al. (2015) showed in simulations that the scalp distribution recovered by average reference is montage dependent in caps with less than 120 electrodes, and that a reference method called “reference electrode standardization technique”, REST (Dong et al., 2017; Yao, 2001), corrects these distortions for lower density caps. REST calculates a source solution for the EEG data set and then adjusts the predicted scalp distribution to an infinity reference. The accuracy of the resulting scalp distribution does not depend on the accuracy of the source solution. We therefore used REST as a reference so that our results would be comparable to results with high density caps using average reference. Results of the mass univariate analysis for average reference were very similar although the N200 training effect (see below) was significant consistently as a positive difference at the mastoids and nearby electrodes and negative at fewer central electrodes (Figures S7, S8, S9). Results using average mastoids were also qualitatively similar: the N200 was more broadly distributed than the other methods (see also Du, Hu, & Fang, 2013; Du, Zhang, & Zhang, 2014), likely due to opposite effects at the mastoids (Figure S4), and the selection negativity was much smaller over the back of the head, likely because the effect extended anteriorly to the mastoids (Figures S2, S3), but still reached significance in some time windows (data available on request).

## Analysis

Based on a previous study with a very similar design and stimulus set, we expected that both training effects (N250) and effects of target features (SN) would be observed at postero-lateral electrode sites. N170 and N250 were measured at electrodes 41 (theta/phi: 92,-45) and 45 (theta/phi: −92,45) where the N170 was largest. N170 was measured as the peak amplitude between 120 and 200 ms. N250 was measured as the mean amplitude between 270 and 300 ms based on previous results with the same stimuli (Folstein, Monfared, et al., 2017). For the N250, our ROI was slightly more narrow than our previous study, which also included channels surrounding 41 and 45. We did this to maximally separate it from the centrally distributed N200/N320 complex. N170 and N250 components were analyzed in 4 × 2 × 2 × 2 repeated measures ANOVAs with factors of Target features (1 to 4 target features), Training (Trained vs. Untrained stimulus set), SOA (fast vs. slow SOA), and Hemisphere (left vs. right).

Because unexpected effects of training were observed at centro-parietal electrodes, we used a mass univariate approach to quantify the timecourse and scalp distribution of attention and training effects (Groppe, Urbach, & Kutas, 2011). All mass univariate comparisons were corrected using False Discovery Rate (Benjamini & Hochberg, 1995). Difference waves between ERPs elicited by conditions of interest were created and evaluated at all electrodes in 10 ms timewindows from 100 to 500 ms. For the parametric effect of attention, which had more than two conditions, a line was fit to the function relating number of target features in non-targets (one, two, three, and four target features) to voltage for each time point in each electrode (voltage = at+c; where a is the slope parameter, quantifying microvolts per target feature, t is number of target features, and c is the y-intercept). Only the slope parameter was used, resulting in a timecourse of linear attention effects at each electrode for each participant (subsequently, **attention effect timecourse, AET**) that was treated as a difference wave in all analyses^1^ (see Figure S10 for further illustration of the AET).

As a supplementary analysis, the onset latency of the SN was compared between trained and untrained stimuli at each SOA using the jackknife procedure, in which latency is measured in each of 16 grand averages with a single participant left out. For each cell in the 2 × 2 (Training x SOA) design, 50% fractional peak latency of the AET between 100 and 400 ms was measured at electrode pair 41 (theta/phi: 92,-45) and 45 (theta/phi: −92,45. The cells were entered into a 2×2 ANOVA and *F* values were corrected for the jackknife procedure according to the equation *F_corrected_ = F*/(*n* −1)^2^ (Kiesel, Miller, Jolicœur, & Brisson, 2008; Ulrich & Miller, 2001).

### Source localization

Neural sources of grand average ERP difference waves were estimated using LORETA-Key software (Pascual-Marqui, 2002), which provides low resolution minimum norm neural sources for EEG. A single set of electrode coordinates was transformed to MNI space using modified functions from EEGLAB, warped to LORETA’s standardized head model using fiducial landmarks. The resulting coordinates were then used to create a transformation matrix using eLORETA, which provides an estimated inverse solution (Pascual-Marqui, 2009). Grand average difference waves between conditions of interest were then transformed to voxel space (sLORETA objects). Finally, current densities for the AET and the difference wave for the trained vs. untrained condition (averaged over all other conditions) were separately averaged over time windows of interest determined by the results of the ERP analyses. This approach of localizing difference waves has been previously used to localize the mismatch negativity (Gil-da-Costa, Stoner, Fung, & Albright, 2013; Takahashi et al., 2013) and ERPs related to object recognition (Schendan & Ganis, 2015; Schendan & Lucia, 2010), and has been validated in a target detection task using combined ERP and fMRI (Mulert et al., 2004). No statistical comparisons were made, as we sought only to identify the strongest neural sources of our effects.

## Results

### Behavioral measures

#### Categorization training

Prototypes elicited the fastest and most accurate responses, followed by Typical and then Near Boundary stimuli, which elicited the slowest and least accurate responses.

Accuracy and reaction time improved with training and the difference between the Typicality conditions decreased with training, also for both measures (Figure 3). To quantify these effects, ANOVAs were conducted with factors of Typicality (prototype, typical, near boundary) and Training Day (days one through six). For both dependent measures, main effects of Typicality and Training Day were significant and the interaction between the two factors was also significant for both measures (Table 1).

**Figure 3.**
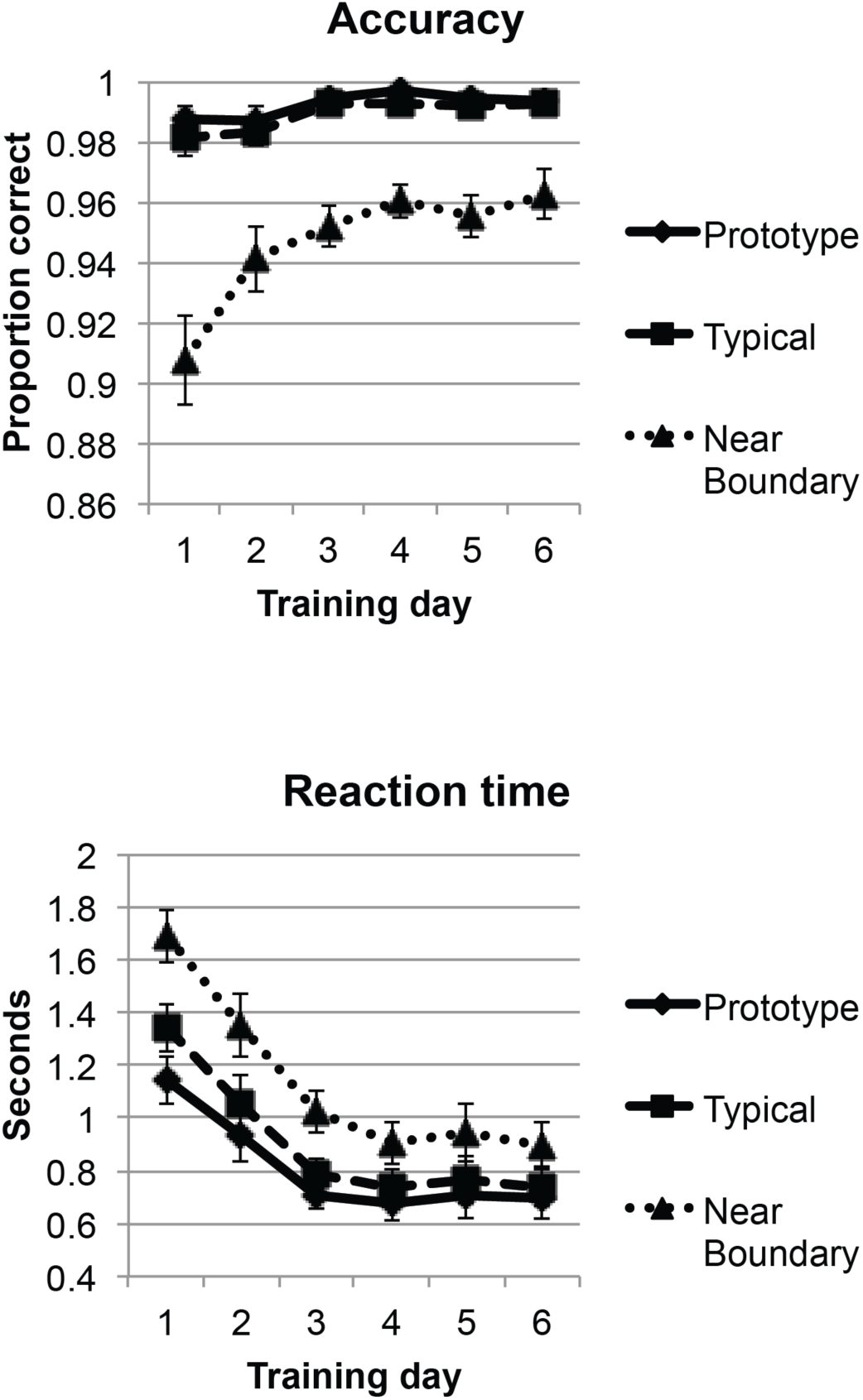
Accuracy and reaction time for each day of categorization training.

**Table 1.** Statistical outcomes for categorization training

| | *F* | *df*(effect,error) | *p* | $\varepsilon$ | $\eta_p^2$ |
| --- | --- | --- | --- | --- | --- |
| *Accuracy* |  |  |  |  |  |
| Typicality | 56.7 | 2,30 | < .0005 | 0.558 | 0.791 |
| Training day | 7.1 | 5,75 | 0.002 | 0.467 | 0.321 |
| Typicality X Training day | 7.6 | 10,150 | < .0005 | 0.339 | 0.337 |
| *Reaction time* |  |  |  |  |  |
| Typicality | 106.5 | 2,30 | < .0005 | 0.553 | 0.542 |
| Training day | 17.8 | 5,75 | < .0005 | 0.698 | 0.876 |
| Typicality X Training day | 26.6 | 10,150 | < .0005 | 0.386 | 0.639 |

#### EEG target detection task

Target and nontarget conditions were analyzed separately as they required different responses (go vs. no-go). Hit rates to targets were quite high (*M =* 94.5). An ANOVA on target accuracy with factors of Training and SOA, yielded no significant main effects or interactions (*F*s(1,15) < 1.5). The same ANOVA conducted on reaction time yielded a main effect of SOA, with faster responses in the fast than the slow SOA conditions (*F*(1,15) = 18.8, *p* < .005, *η*^2^_p_ = .556). No other effects reached significance.

False alarms to nontargets were highest to stimuli with four target features, which were the most similar to the target, and higher for the fast than slow SOA (Figure 4), suggesting that the faster SOA increased the difficulty of the task. Accuracy for nontargets was assessed in a 5 × 2 × 2 ANOVA with factors of Target features (zero through five), Training, and SOA. Consistent with the pattern in Figure 4, the ANOVA yielded main effects of Target features (*F*(4,60) = 51.0, *p* < .0005, ε = .286, *η*^2^_p_ = .773) and SOA (*F*(1,15) = 26.6, *p* < .0005, *η*^2^_p_ = .640). The Training x Target features interaction, indicating a larger decrement in accuracy non-targets with four target features compared to other non-targets, approached significance (*F*(4,60) = 2.96, *p* = .072, ε = .467, *η*^2^_p_ = .165).

**Figure 4.**
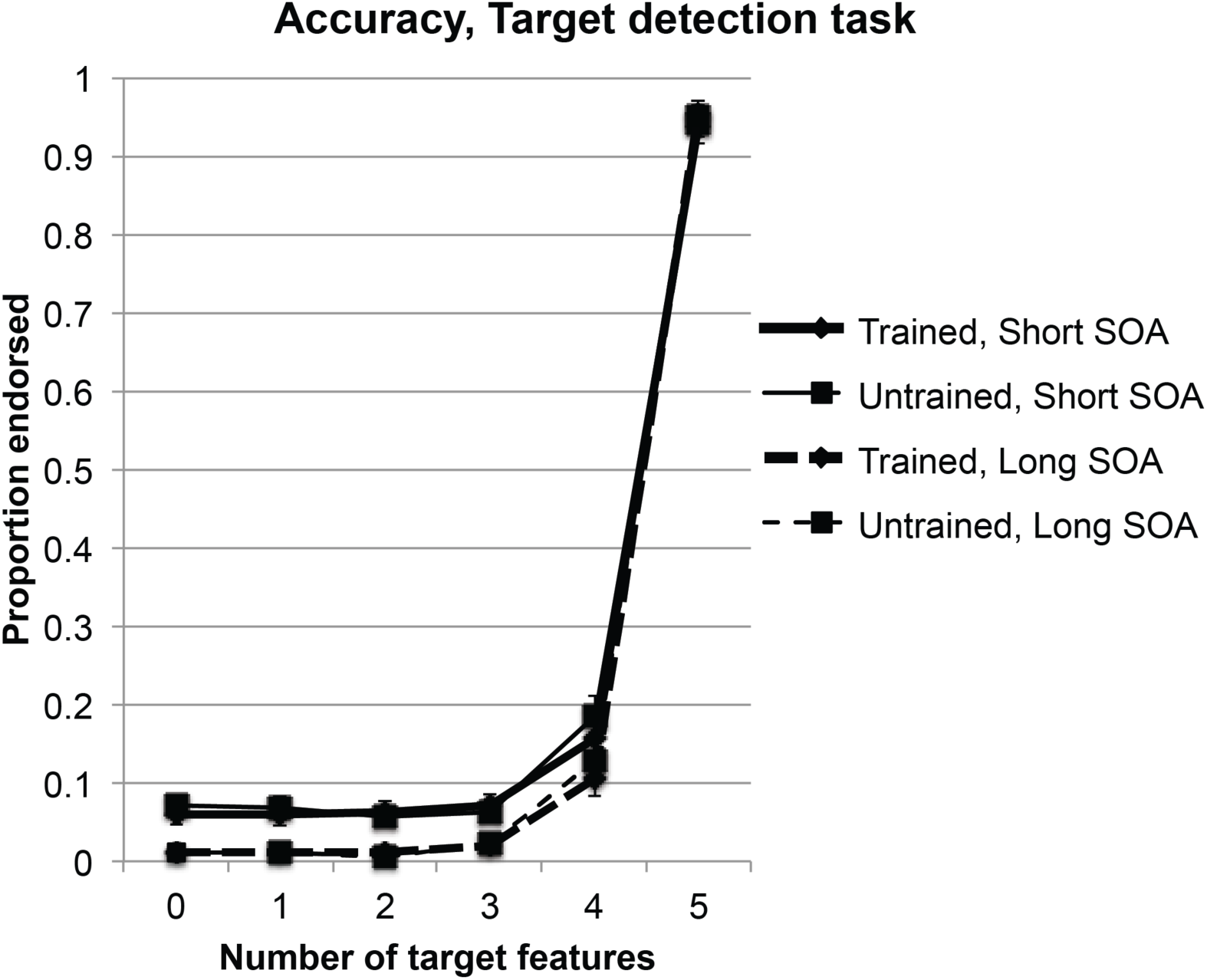
Proportion of each stimulus type endorsed as targets during the EEG target detection task. X-axis shows number of target features in each condition, including targets, which had five target features.

## Event related potentials

### Component of interest analysis

Prior to the experiment, we hypothesized that the N170 and the N250 would be sensitive to training. We therefore analyzed these components at electrodes or interest.

#### N170

The N170 was more negative in the slower SOA condition. The repeated measures ANOVA with factors of Target features, Training, SOA, and Hemisphere returned only significant effects of SOA (*F*(1,15) = 33.1, *p* < .001, *η*^2^_p_ = .688) and an SOA x Hemisphere interaction (*F*(1,15) = 8.09, *p* < .05, *η*^2^_p_ = .350) reflecting a larger effect of SOA in the left than the right hemisphere.

#### N250 and SN

Trained stimuli were slightly more negative than untrained stimuli at postero-lateral electrodes (270 to 300 ms), but the effect was quite small (Figure 5). In contrast, the SN was robust at the same electrodes (Figure 6). A repeated measures ANOVA with the same factors as the N170 was applied to the N250 measurement. As expected, there was a large main effect of Target features (*F*(3,45) = 37.1, *p* < .0005, *ε* = .443, *η*^2^_p_ = .716), an effect we interpret as the SN. As in our previous study (Folstein, Monfared, et al., 2017), the interaction between Target features and Training did not approach significance (*F*(3,45) < 1), but, in contrast to our earlier study, the main effect of Training was not significant either (*F*(1,15) = 1.01, *p* = .33, *η*^2^_p_ = .063). Interestingly, the interaction between Training and SOA approached significance (*F*(1,15) = 4.07, *p* = .06, *η*^2^_p_ = .213), reflecting a larger effect of Training for the longer SOA. Exploratory follow-up ANOVAs were performed at each SOA with the same factors as before, but the effect of Training was not significant for either analysis.

**Figure 5.**
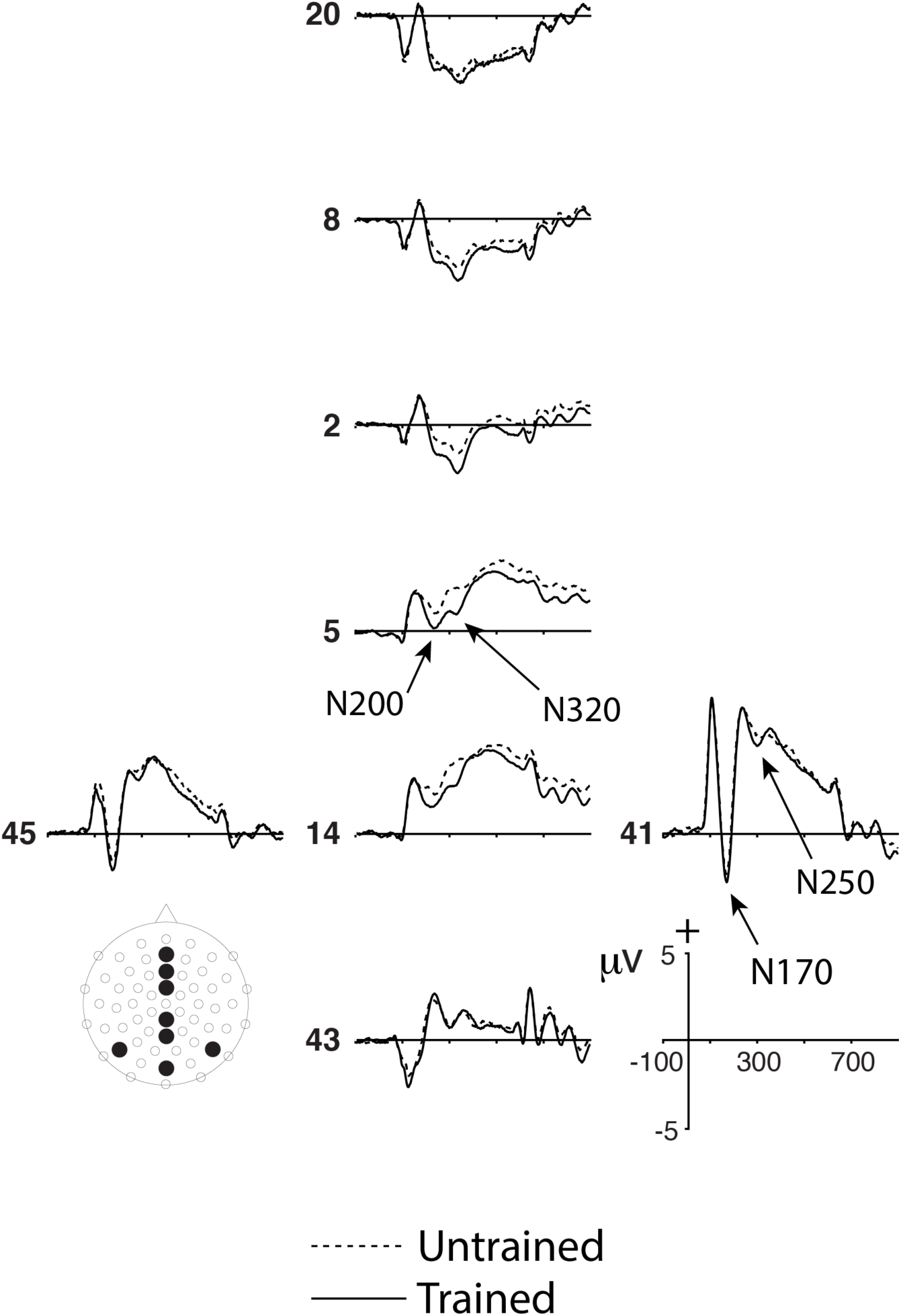
ERPs for trained and untrained stimulus conditions, collapsed over all other factors. The N200 and N320 components were most prominent over centro-parietal and central electrodes. Small N170 and N250 training effects are apparent at postero-lateral sites, but these effects did not reach significance.

**Figure 6.**
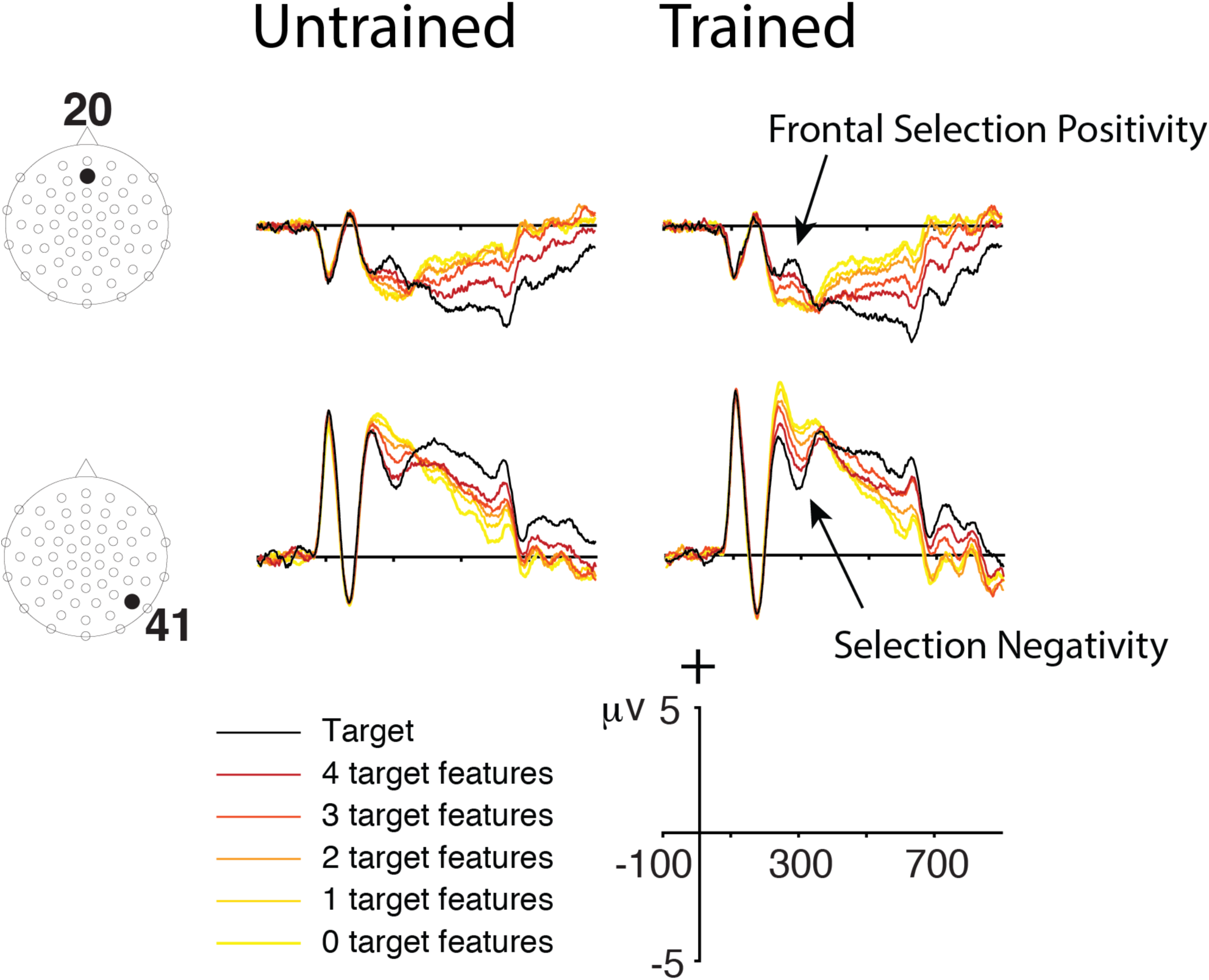
Cumulative selection negativity (SN) and frontal selection positivity (FSP) at representative frontal and postero-lateral electrodes for trained (right) and untrained (left) stimuli. The cumulative SN becomes increasingly negative in stimuli with more target features as SNs from more and more target features summate.

### Mass univariate analysis

Mass univariate analysis was performed to explore the effect of training and compare its timecourse and scalp distribution to the SN and FSP.

#### Selection Negativity and Frontal Selection Positivity

As in our previous studies, we found robust cumulative SN and FSP effects with increasing numbers of target features (Figure 6). Figure 7 shows the timecourse of the linear effect, or attention effect time course (AET, see Methods). Inspection of Figure 7 revealed that the onset of the SN was earlier for the trained than the untrained stimuli. We directly compared the onset latency of the trained and untrained SN using the jackknife procedure (see Methods). Fractional (50%) peak latency was entered into a 2 × 2 × 2 ANOVA with factors of Training, SOA, and Hemisphere (electrode 41 vs. 45). Only the effect of Training reached significance (*F*_corrected_(1,15) = 9.21, *p* < .01), reflecting the earlier onset of the SN to the trained stimuli.

**Figure 7.**
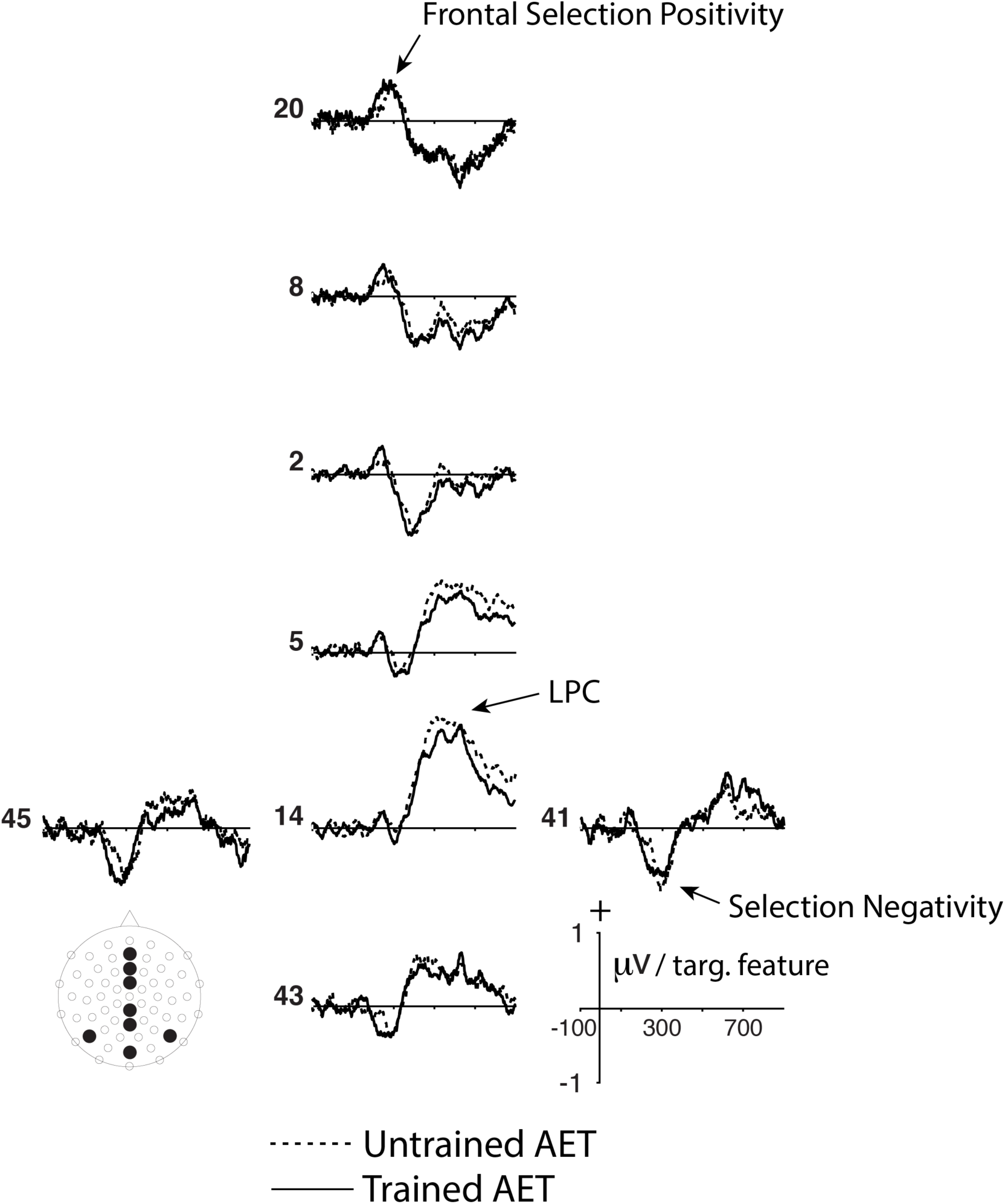
The SN portion of the attention effect timecourse (AET) onset earlier for the trained than untrained stimuli. Each sample in the traces was generated by fitting a line to the effect of target features on voltage (conditions with 1 through 4 target features were used). The selection negativity portion of the AET is negative because ERPs were increasingly more negative in stimuli with more target features. This approach allowed us to analyze the timecourse of the effect and facilitated mass univariate analysis. See Figure S10 for further illustration.

We then conducted a mass univariate analysis to explore interactions between attention and the other factors of the design. The AET did not differ between SOAs and no interaction was found at any timepoint between the factors of Training and SOA^2^ and we therefore collapsed the AET over SOA conditions. To explore the interaction between the effect of training and the effect of attention, we compared the AET elicited by the trained stimuli with the AET elicited by the untrained stimuli in the mass univariate analysis. No electrodes were significant at any timepoint, demonstrating that there were no strong interactions between training and attention.

To compare the timecourses of the trained and untrained AET in greater detail, we conducted mass univariate analyses on each condition. The effect of attention was negative in large clusters of posterior and postero-lateral electrodes and positive in large clusters of frontal electrodes during the SN/FSP epoch for both the trained and untrained stimuli (Figure 7, 8). The SN in the trained condition (Figure S2) reached significance at about 190 ms, 60 ms earlier than the untrained condition, which reached significance at about 250 ms (Figure S3). The SN elicited by the trained stimuli was also sustained for a longer period, about 120 ms at posterolateral electrodes compared to about 60 ms to 80 ms for the untrained condition. The FSP showed a similar pattern, but was more difficult to quantify, as it tended to shift scalp distribution more quickly, moving from fronto-central to more fronto-polar areas.

#### Later components

A second complex of components sensitive to target features arose after the resolution of the SN/FSP portion of the AET. These components included a frontally distributed negative component, most prominent for the trained stimuli, that was significant at frontal electrodes starting at about 360ms in the trained condition (Figure 7, S2, S3) and overlapped with a posterior P300. The LPC reached significance somewhat later between 390 and 410 ms and remained significant to the end of the analysis window at 500 ms.

#### N200 and N320

The main effect of training was tested in the mass univariate analysis by averaging across all SOA and target feature conditions to create ERPs for the trained and untrained conditions^3^ (Figure 5). This contrast revealed two time windows in which large numbers of channels were significantly more negative to trained than untrained stimuli, one between 200 and 250 ms and a second between 290 and 370 ms. Rather than a postero-lateral scalp distribution, as would be expected for the N250, both time windows had central and centro-parietal scalp distributions (Figure 8, Figure S4). We will refer to the first as the N200 and the second as the N320. In both clusters, trained stimuli were significantly more negative than untrained stimuli at central and centro-parietal channels, with the opposite effect at some lateral and lower eye channels (Figure S4).

**Figure 8.**
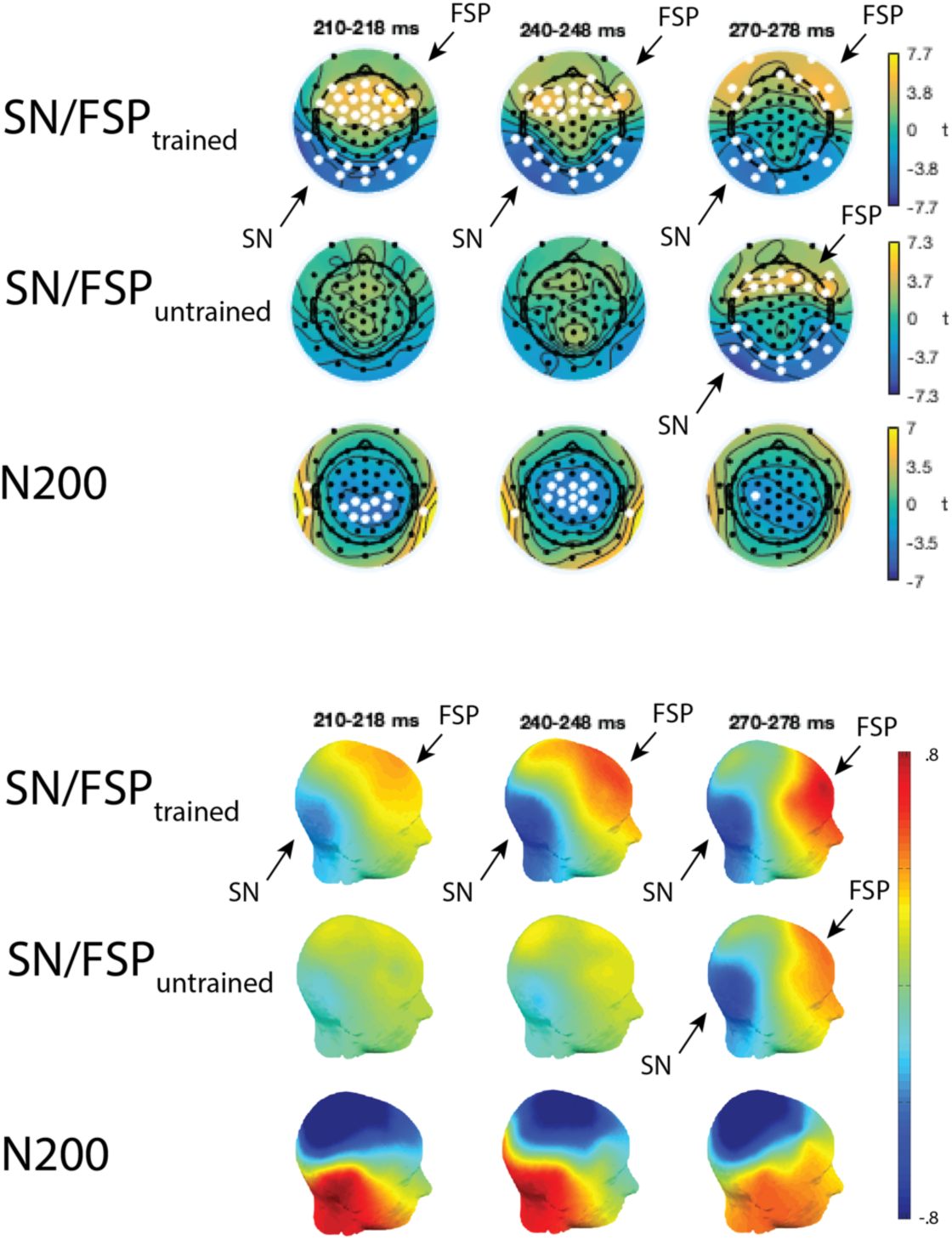
Top: topographical maps from three representative epochs of the mass univariate analysis (see Figures S2 through S5 for full mass univariate results). Col or scale corresponds to t-values and white electrode locations represent electrodes with significant differences within each time window. The first two rows show electrodes where the AET was significantly different from zero, reflecting the main effect of target features. The effect onsets earlier for trained (first row) than untrained (second row) stimuli. The N200 (third row) had a similar time course to the SN in the trained condition. Bottom: topographical voltage maps of the three effects.

To rule out the possibility that the N200 might be an enhanced no-go N2 for trained stimuli, we repeated the analysis but this time restricted the analysis to targets, which required a “go” response. Trained stimuli were again more negative than untrained stimuli at centro-parietal electrodes during the same time window from 210 to 260 milliseconds. The effect was weaker and more right lateralized than the original analysis, reaching significance at fewer electrodes (Figure S5).

The N320, measured at the original four non-target conditions, also had a centro-parietal distribution at its onset at about 280 ms, but again had a central distribution at its peak. The effect of training ended at 370 ms.

Finally, it should be noted that some electrodes just dorsal to the postero-lateral channels where the N250 was predicted did reach significance during the N250 time window (Figure S4). The overall scalp distribution during this time range is clearly centro-parietal, however (Figure S6).

#### Comparison of N250 and N200

The overall pattern of training effects observed so far was rather clear: ERPs to trained stimuli were more negative than ERPs to untrained stimuli at central and centro-parietal channels, whereas no effect of training was observed at postero-lateral channels in the N170 or N250 epoch, neither in mass univariate analysis nor at hypothesis driven channels of interest. Nevertheless, the effect of training at poster-lateral channels trended in the expected direction, with trained stimuli more negative than untrained stimuli. To compare the two components directly, we first measured the N200 at centro-parietal electrodes 1, 4, 5, 6, 13, 14, 15 from 200 to 270 ms. We then conducted 2×2 ANOVAs with factors of Component and Training separately for N250 (again measured from 270 to 300 ms) in each hemisphere, averaging across all other conditions. Main effects of Component and Training were significant for both hemispheres (*F*s(1,15) > 6, *p*s < .05), but the interaction was not significant for either hemisphere (*F*s(1,15) < 1). This may not be surprising given that the centrally distributed N200 and N320 likely influenced the postero-lateral electrodes.

### Source Localization

Figure 9 shows the current source density for the grand average trained AET (SN) from 190-248 ms and 250-318ms, the untrained AET (SN) from 250-340 ms (where is was significant), the N200 from 190-248 ms, and the N320 from 290-370 ms. Note that the figures are scaled to 75% maximum to emphasize the local maxima of current density in a given condition rather than absolute differences between conditions. Whereas both the SN localized to the ventral surface of the occipital and temporal cortex, the largest SN sources were in the right hemisphere while the largest N200 and N320 sources were in the left hemisphere. Because this analysis was exploratory, we do not attempt to characterize differences between conditions in further detail.

**Figure 9.**
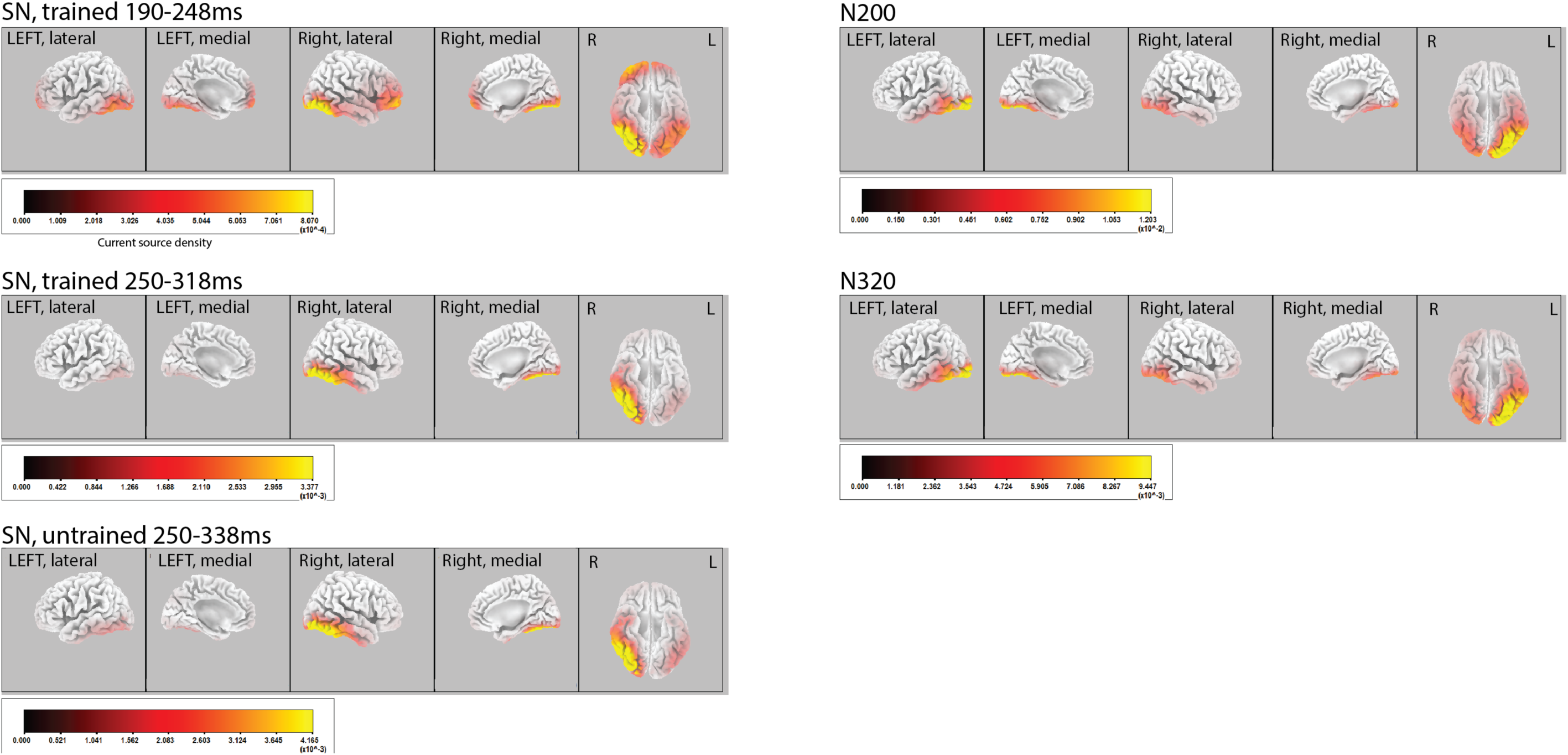
Sources of the SN/FSP, N200, and N320. The AET and the difference wave between trained and untrained stimuli were localized using eLORETA. Scale maximums are 75% maximum current density for each condition. Both the SN/FSP and the N200/N320 had strong neural sources in ventral temporal areas, but the strongest sources for the SN/FSP were in the right hemisphere and the strongest N200/N320 sources were in the left hemisphere.

## Discussion

Using a target detection paradigm with ERPs as our dependent measure, we tested the effect of training on the neural correlates of attentional feature selection. This in turn allowed us to isolate effects that were independent of attentional feature selection. We found that 1) the onset of attentional selection (the postero-lateral SN) was 60 ms earlier for trained than untrained features 2) separate effects of training, the N200 and N320, were observed that did not interact with attention and had a distinct centro-parietal scalp distribution, consistent with previously observed effects related to orthography, and highly inconsistent with previously observed perceptual expertise effects 3) source localization of difference waves revealed local maxima in the right ventral temporal cortex for the SN and in the left ventral temporal cortex for the N200.

### Effect of training on attention

In the current study we sought to answer the question of whether category learning altered the neural correlates of attentional feature selection, which we operationally defined as the SN/FSP. After six sessions of category learning, attentional selection of the trained features was earlier and somewhat more sustained than selection of untrained features. When measured near its peak (270-300 ms), the SN did not differ significantly between the trained and untrained stimuli, nor did the AET differ between trained and untrained stimuli when compared in the mass univariate analysis. This suggests that, as in our previous study, training had little effect on the peak amplitude of the selection negativity. Category learning did affect the latency of the SN, however, reducing it by about 60 milliseconds. Furthermore, the mass univariate analysis showed that the offset of the trained SN was only slightly earlier than offset of the untrained SN such that the duration of trained SN was about 130 ms compared to about 90 ms for the untrained SN. To our knowledge, ours is the first study to demonstrate that category learning alters the latency of attentional feature selection.

Our results demonstrate that category learning results in facilitation of attentional processing that generalizes to other tasks. The earlier onset, more sustained SN to trained stimuli was quite similar to the result of (Smid et al., 1999), who observed an earlier onset SN to salient, easily perceived target features than smaller, less salient visual shape features. This suggests that the attention effect we observed was driven by an increased ability to perceptually discriminate between the features of the trained stimuli and links our result with previous demonstrations of task independent increases in discriminability resulting from category learning (Folstein, Palmeri, Van Gulick, & Gauthier, 2015). Thus, perceptual training did alter attentional feature selection, but probably by virtue of changes to perceptual representations (c.f. Dieciuc, Roque, & Folstein, 2017).

Several later components were also modulated by target features, including the LPC, consistent with several previous studies (Azizian, Freitas, Watson, & Squires, 2006; Dieciuc et al., 2017; Folstein, Fuller, et al., 2017; Folstein, Monfared, et al., 2017). These components are not addressed by our hypotheses and, based on the mass univariate analysis, there was no evidence that the AET differed greatly between trained and untrained stimuli during this time range.

### The effect of training

Based on many previous studies (Jones et al., 2018; Scott et al., 2006, 2008), including a training study with the same stimuli but briefer training period (Folstein, Monfared, et al., 2017), we expected trained stimuli to elicit higher amplitude N170 and N250 components at postero-lateral electrode cites than untrained stimuli. In the current study, both the postero-lateral N170 and the N250 trended in the predicted direction, but the effect of training did not reach significance for either component. A much larger effect of training was observed centered at central or centro-parietal channels, at which there were two discernable peaks, one at 230 ms and a second at 328 ms. Mass univariate analysis suggested two separate epochs in which large numbers of central and centro-parietal electrodes reached significance, one from 190 to 250 ms and the second from 290 to 370 ms. These effects not only had an unexpected scalp distribution, but also had unexpected time courses. The N200 was too early and the N320 too late to match the usual N250 (230-330 ms (Scott et al., 2006, 2008); 210-330 ms (Jones et al., 2018); 270-300 ms (Folstein, Monfared, et al., 2017)).

Effects related to general recognition or familiarity can also be ruled out. Across many studies, stimuli that are more familiar (Curran & Cleary, 2003; Folstein & Van Petten, 2004; Schendan & Kutas, 2003) are more positive at central electrode channels in similar time windows than less familiar or unrecognized stimuli. Similarly, stimuli that are easier to identify due to cannonical viewing angle (Schendan et al., 2003) or reduced noise (e.g. Schendan & Maher, 2009) are more positive at central and frontal electrode cites than more difficult control conditions. When comparing more vs. less impoverished line drawings of real (but not impossible) objects (Schendan et al., 2015) found a postero-lateral N250 component very similar to N250 components observed for objects of expertise in training studies. Contrary to what would be predicted based on these studies, our trained stimuli elicited more negative ERPs than untrained stimuli at centro-parietal electrode channels.

More similar findings to our own are reported by studies of orthography. Ruz et al. (2008) reported an N200 greater to orthographic judgments than phonological and semantic judgments from 190 to 260 ms. The component had a left lateralized posterior distribution, but appears to have extended dorsally and medially relative to the earlier N170 observed in the same study. Bentin, Mouchetant-Rostaing, Giard, Echallier, and Pernier (1999) observed an N320 component from 270 to 370 ms, quite similar to the timecourse of the N320 component we observed, was more negative to pronounceable than non-pronounceable stimuli over a broad range of scalp locations, including frontal and central locations.

Studies in which subjects are trained in artificial languages have also sometimes found effects in time windows similar to the N200/N320. McCandliss, Posner, and Givon (1997) identified an unusually late word-sensitive “N1” time window from 170 to 230 ms and a second time window from 280 to 360 ms. The same study found that pseudowords trained to be associated with novel meanings (“Keki” language) elicited more negative ERPs than control pseudowords in the second time window, similar to the N320 (scalp distribution was not analyzed in detail). Yoncheva, Wise, and McCandliss (2015) found that the scalp distribution a component labeled the N170 was affected by the type of training participants received on an artificial script, being more left lateralized after orthographic training than logographic training. Importantly the effect was from 197-222 ms. A similar study (Yoncheva, Blau, Maurer, & McCandliss, 2010) found effects from 186-190ms, suggesting that orthographic training effects can be later than the usual N170.

Other studies have used masked priming to reveal word-level orthographic representations with the reasoning that ERPs related to lexical accessed will be reduced by masked priming, which is predicted to increase the fluency with which representations are accessed. A central or centro-parietal N250 component between 200 and about 300 ms is reduced when words are primed by the same word (different case) compared to different words (Holcomb & Grainger, 2007) and with transposed letters (Grainger et al., 2006) compared to the word with the same number of letters replaced, which served as a less word-like control. A similar reduced N250 is observed when words are primed by consonant-only versions of the same word vs. vowel-only versions, the former having been shown to be a more effective orthographic prime (Carreiras et al., 2009).

A component very similar to the N200 has also been observed in studies of Chinese orthography in which participants perform an unrelated task (e.g. lexical decision) on sequences of two visible Chinese words. A centro-parietal N200 component, peaking at about 220 milliseconds and measured from about 190 to 235 ms (e.g. Du et al., 2013; Zhou et al., 2016), is increased when the same word is presented twice, but not when the target word is preceded by an orthographically unrelated homophone (Zhang et al., 2012) or a semantically related word (Du et al., 2014). The effect also appears limited the Chinese words as opposed to non-native languages such as Korean (Zhang et al., 2012). Overall, the N200 is interpreted to be sensitive to orthographic visual processing of Chinese characters. The similarity of this component to the N250r (Schweinberger & Neumann, 2016) is striking as both components are sensitive to immediate and slightly delayed repetitions and are sensitive to expert object domains, the N250r to faces and the N200 to words. Unfortunately, no training studies have tested the effect of training on the N250r and it is also unknown whether objects of real-world expertise, such as birds or cars, elicit an enhanced N250r in experts.

Finally, source localization of the N200 difference wave revealed a strong local maximum in left ventral temporal cortex. Although preliminary, this result is also consistent with an orthographically linked neural source, such as the visual word form area (VWFA), located in the left fusiform gyrus (Dehaene et al., 2011). Future studies will be needed to determine if this is the case but, given that our stimuli are animals, our paradigm could offer important clues about the preconditions for recruiting VWFA, which are currently controversial (Dehaene et al., 2011; Price et al., 2011).

Overall, there is a strong circumstantial case that the N200/N320 components that were elicited by training are related to orthography, suggesting that we might have induced word-like perception of stimuli that were initially perceived as animals. Broadly consistent with this conclusion, a standard postero-lateral N250 training effect was observed in the same stimuli with the same training but with a briefer training period (Folstein, Monfared, et al., 2017). This suggests that our stimuli are processed as objects early in learning and may only transition to a new kind of representation, perhaps orthography, later.

Three possibilities stand out as explanations for recruitment of ERP effects that differed from the standard N170/N250 training effects that have been observed by others. First is our rule-based categorization training, which called attention to individual features in the exemplars, might have encouraged processing the stimuli in a part-based manner, which is associated with left hemisphere representation (Yovel, Yovel, & Levy, 2001). Second, the interchangable nature of our stimulus features might have encouraged part-based processing, even though all features had to be attended to successfully apply the rule. Third, the stimuli were defined by black outlines and therefore contained many line junctions, to which the VWFA is known to be sensitive (Dehaene et al., 2011).

What we can say is that our training paradigm did not explicitly encourage mappings between the stimuli and language related representations such as phonemes or meaningful words (Maurer, Blau, Yoncheva, & McCandliss, 2010; McCandliss et al., 1997; Mei et al., 2013; Moore, Durisko, Perfetti, & Fiez, 2014; Song, Hu, Li, Li, & Liu, 2010; Xue et al., 2006; Yoncheva et al., 2010; Yoncheva et al., 2015). Rather, our task was a simple naming task similar to previous training studies with objects. Also, although our stimuli had lines and even look a bit like hieroglyphics in retrospect, it is clear that they are not an artificial script, pseudofont, pseudoword, or foreign script as the stimuli were in most of the artificial language studies cited above, with the notable exceptions of Moore et al. (2014), who mapped faces onto phonemes and Song et al. (2010), who mapped geometric stimuli onto words, both recruiting the left fusiform. Thus, while further experiments will be necessary to determine what properties of our training or stimuli caused recruitment of the N200/N320, our stimuli and paradigm could offer an intriguing opportunity for identifying the preconditions for recruiting orthography-linked brain areas.

### Relationship between training and attention

The main purpose of the current study was to tease apart the effect of training on neural correlates of attention from other effects of training. One motivation for this attempt was studies evaluating expertise effects using a match-to-category task. We argued previously (Folstein, Monfared, et al., 2017) that experts performing the match to category task could use a different attentional strategy than novices, such that the training related N250 could be driven by differences in attention to the stimulus features, especially given its late onset and similarity to the SN. Our parametric manipulation allowed a nuanced comparison between attentional feature selection processes in trained and untrained stimuli and identification of components that were insensitive to feature selection. Not only were the N200 and N320 insensitive to the number of relevant features in the stimulus, they had a completely different scalp distribution, a departure from our previous study, in which the time course and scalp distribution of training and attention effects overlapped, but were additive. Furthermore, their neural sources were strongest on the left, while the neural sources of the attention effect were strongest on the right.

Whereas, the N200 and N320 components were not sensitive to the same strategic feature selection processes as the SN, their late time windows suggest that they are driven by feedback (e.g. Ganis et al., 2007; Lamme & Roelfsema, 2000; Schendan et al., 2015). Price et al. (2011) proposed that learning to recognize words (or, by generalization, other types of stimuli) results in automatic generation of predictions carried by feedback signals to perceptual cortex (e.g. Friston & Kiebel, 2009). With more prediction comes greater prediction error and therefore increased cortical activation. This activation was predicted to be lowest for novices, who generate no predictions, highest early in learning, and somewhat lower after longer term learning, because more accurate predictions result in less prediction error. Interestingly, the N250 is commonly observed after training studies lasting six to ten days but is observed less consistently in real-world experts. A recent report found a decreased N250 in other race faces compared same race faces, even though same race faces are associated with greater expertise, and also reported a negative correlation between N250 amplitude and car expertise (Herzmann, 2016). Taken together, these findings are consistent with the prediction that intermediate levels of experience result in the strongest feedback. The current results match the further prediction that this feedback is not strategic.

A final intriguing finding was that the N200 training effect coincided very closely in time with the latency decrease observed for the trained SN, both onsetting at about 190 ms. Even though the differing scalp distributions, source localization, and functional selectivity strongly suggested that the two effects had separate neural sources, this temporal convergence suggests that they could be causally connected in some way. Further studies will be needed to explore this possible relationship.

## Conclusion

The purpose of this study was to examine the effect of categorization training on attentional modulation and attention-independent neural representations. We found that six days of training affected both types of representations, causing attentional selection of target features to become earlier and more sustained, but also recruiting ERPs that were unaffected by features shared with a target. The two effects had radically different scalp distributions and were localized to separate hemispheres, suggesting that they had different neural correlates. The ERP components elicited by the training were unusual and more consistent with orthographic processing than object processing, a surprising result given that we trained participants to name animal stimuli. Future studies will be required to determine if the key variable in recruiting these components is part-based processing or simple physical features such as line junctions. Overall, our data suggest that training alters neural feedback to perceptual areas related to both strategic and non-strategic processes.

## Acknowledgements

This research was funded by Florida State University.

## SUPPLEMENTAL MATERIALS

### Supplemental Methods

#### Category learning

The study phase began with a review of the five features of the trained set (e.g., “head, body, mane, legs, and tail”) followed by an explanation that each feature had two “types”, Mog features and Vec features. Participants were then shown the categorization rule: “An alien is a Mog if it has at least three Mog features [e.g., Mog head, Mog body, etc.] and a Vec if it has at least three Vec features [e.g., Vec head, Vec body, etc.]”. Next, the participant was shown the Mog prototype and the Vec prototype and told that they would view a number of stimuli of each type (i.e., 16 Mog, 16 Vec). Finally, participants were allowed to view all 16 exemplars in each category at their own pace, one at a time, along with their labels and the features that were diagnostic for that exemplar (e.g. “This is a Mog: it has Mog head, Mog tail, and Mog body”). When all of the exemplars both categories had been viewed, participants were given the option to view the exemplars again or continue to the training phase and return to the study phase later if necessary.

During the training phase, participants categorized unlabeled exemplars of Mogs and Vecs in random order using button presses and received accuracy feedback after each trial. Participants were trained over the course of six sessions, completing a total of 8,800 trials (session 1: 1120 trials; sessions 2 to 6: 1536 trials each). All stimuli were presented with equal frequency. On each trial, a stimulus appeared and remained on the screen until the participant categorized it or 30 seconds elapsed. Accuracy feedback was presented 500 milliseconds after the response (e.g. “Correct, Mog”) for 700 milliseconds, followed by a one second interstimulus interval. The message “Out of time, press C to continue” appeared if participants did not respond in 30 seconds. At the end of each block of 32 exemplars, participants were informed of how many blocks they had completed and offered the opportunity to review the labeled stimulus set.

#### Target detection task

During EEG recording, participants responded to rare target stimulus with a button press, ignoring all other stimuli. Trained stimuli were presented during four runs (360 trials per run) and untrained stimuli were presented during four runs, with stimulus set alternating every two runs. Which set was presented first was counterbalanced across participants. Half of the runs in each training condition used a fast stimulus onset asynchrony (SOA) and half used a slow SOA, alternating every run with order counterbalanced across conditions. The target was always the Mog prototype (recall that the particular stimuli assigned to be the Mog and Vec prototypes were randomized across pairs of participants). Non-target stimuli varied in the number of features they shared with the target. Typical Mogs shared four features with the target, near-boundary Mogs shared three features with the target, near-boundary Vecs shared two features with the target, typical Vecs shared one feature with the target, and the Vec prototype shared zero features with the target. Five out of the ten near-boundary exemplars in each category were pseudo-randomly selected with the constraint that all Mog features and all Vec features were equally frequent in the selected set. This left five exemplars in each non-target condition. Mog prototypes (targets) and Vec prototypes each had one exemplar, presented five times per block. All other stimuli were presented once per block (five presentations per condition), resulting in 30 stimuli per block and a target probability of 16.67%. For the fast SOA condition interstimulus interval was jittered randomly between 800, 900, 1000, and 1100 ms. This was doubled for the slow condition for ISIs of 1600, 1800, 2000, and 2200 ms. A fixation cross appeared in the middle of the screen during each interstimulus interval. Participants received a break every 60 trials, at which point they received a reminder of the target stimulus and instructions to keep their eyes on the location of the fixation cross. Responses between 350 and 2000 ms after the target were counted as correct.

**Figure S1.**
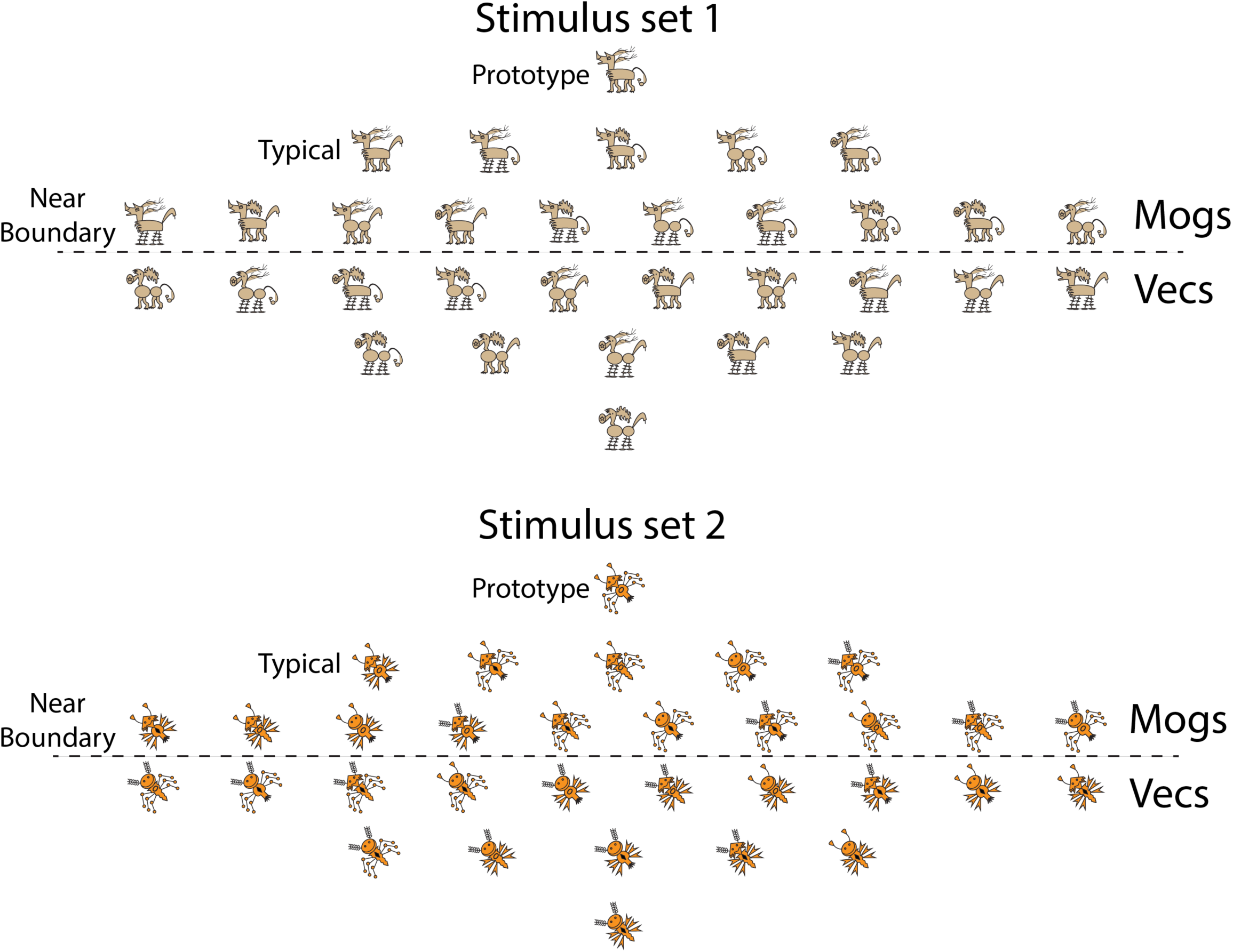
Full set of stimuli used in the experiment. For each yoked pair of participants, stimuli in each set were chosen at random to serve as the prototype. Because of the factorial structure of the stimulus set, each stimulus has an anti-stimulus with which it shares zero features (anti-prototype), five stimuli differing by only one feature (Typical), etc. Thus, assignment of stimuli to conditions was determined by the choice of prototype.

**Figure S2.**
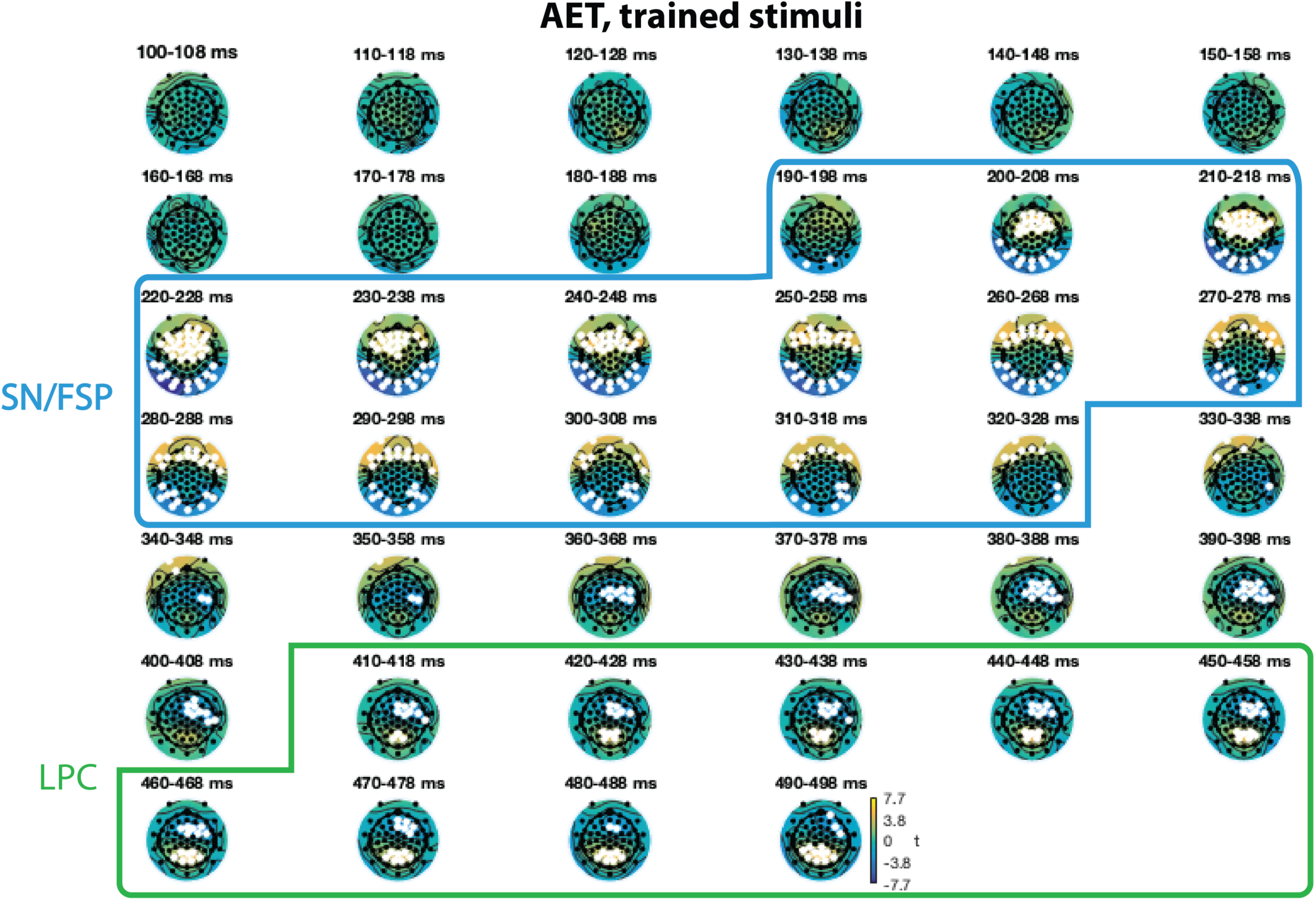
Mass univariate analysis of the trained attention effect timecourse (AET, see methods, Figure S10) in 10ms time windows from 100 to 500 ms. White dots show electrode positions in which the AET significantly differed from zero within each time window. Multiple comparisons were corrected to p < .05 using False Discovery Rate. Blue box shows the approximate temporal boundaries of the SN/FSP component, which is characterized by positive differences at frontal electrodes and negative differences a postero-lateral electrodes. At 330 or 340 ms, the SN resolves into a more centrally distributed negative effect of target features. The green box shows the LPC, characterized by a positive effect at parietal electrodes.

**Figure S3.**
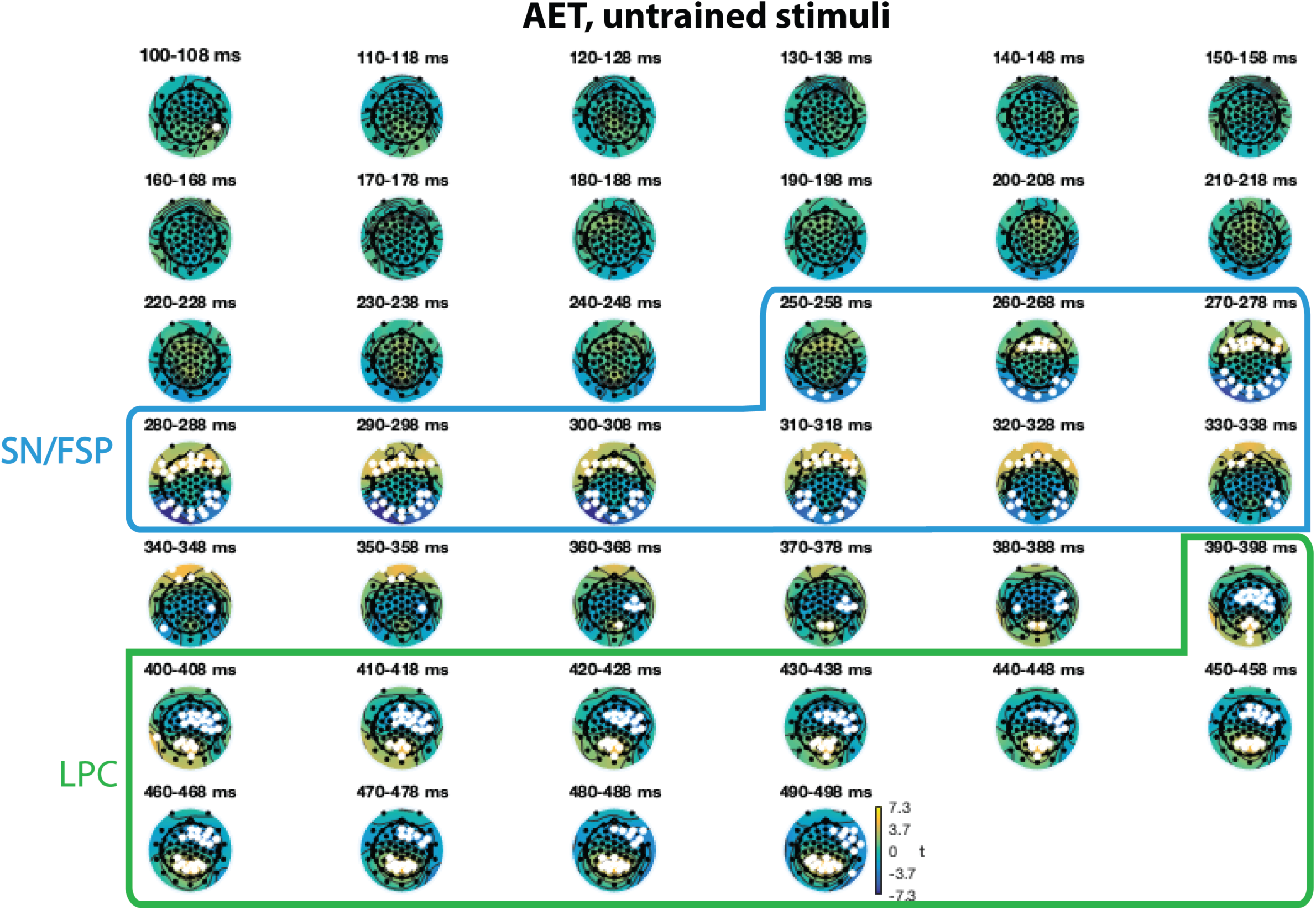
Mass univariate analysis of the untrained AET in 10ms time windows from 100 to 500 ms. White dots show electrode positions in which the AET significantly differed from zero within each time window. Multiple comparisons were corrected to p < .05 using False Discovery Rate. Blue box shows the approximate temporal boundaries of the SN/FSP component, which is characterized by positive differences at frontal electrodes and negative differences a postero-lateral electrodes and resolves between 340 and 350ms. The frontal negative effect becomes prominent at about the same time as the LPC, at about 390 ms, although some posterior parietal electrodes have a positive effect before that time.

**Figure S4.**
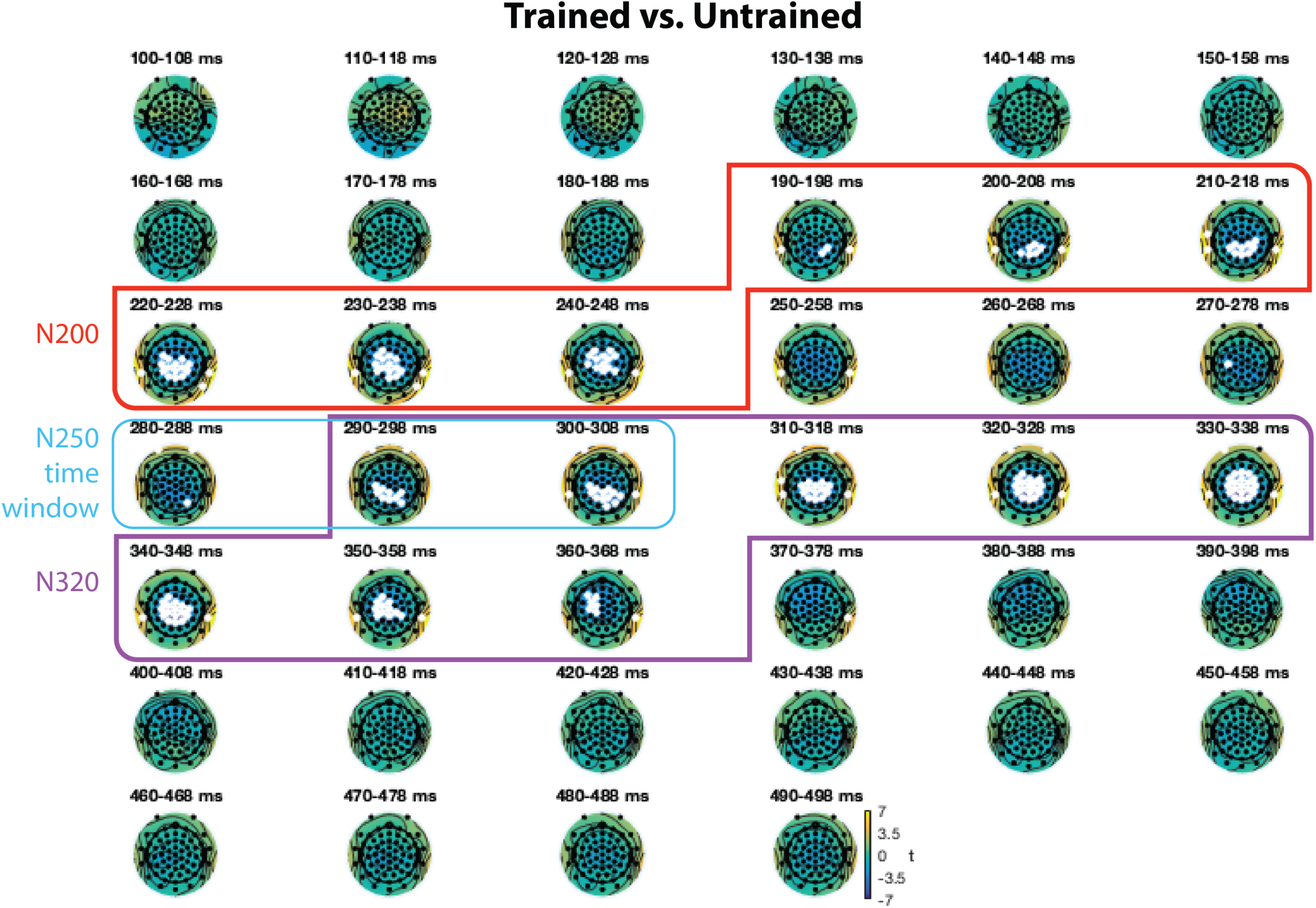
Mass univariate analysis of the trained vs. untrained difference wave for non-targets with 1 through 4 target features. Multiple comparisons were corrected to p < .05 using False Discovery Rate. Red box shows the N200 time window, purple box shows N320 time window. Note that they are punctuated by about 40 ms when very few comparisons are significant. Light blue box shows the a priori time window of the N250. During this window, some more ventral electrodes do become significant, but the overall scalp distribution is centro-parietal (see Figure S6).

**Figure S5.**
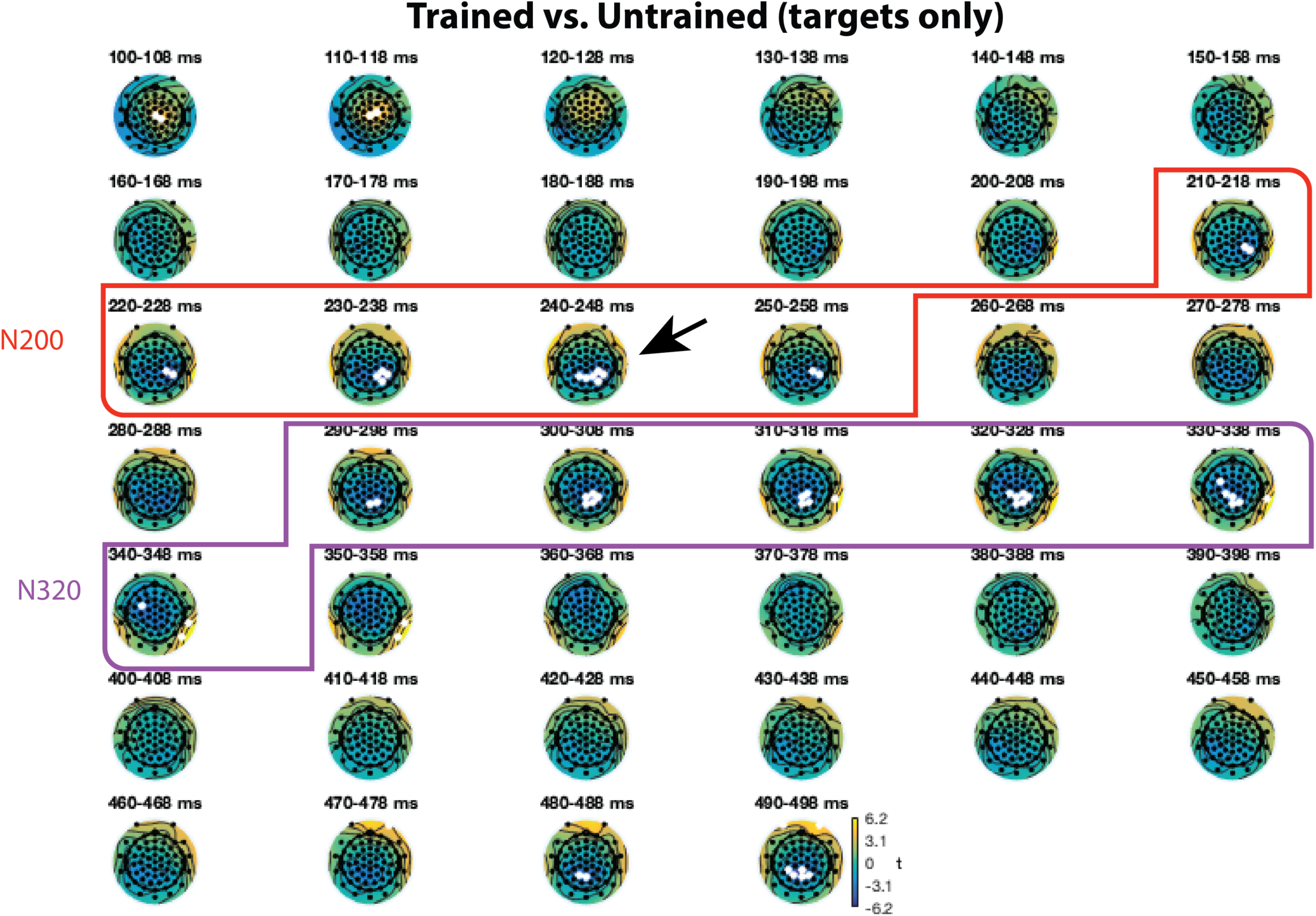
Mass univariate analysis of the trained vs. untrained difference wave for targets only. Multiple comparisons were corrected to p < .05 using False Discovery Rate.

**Figure S6.**
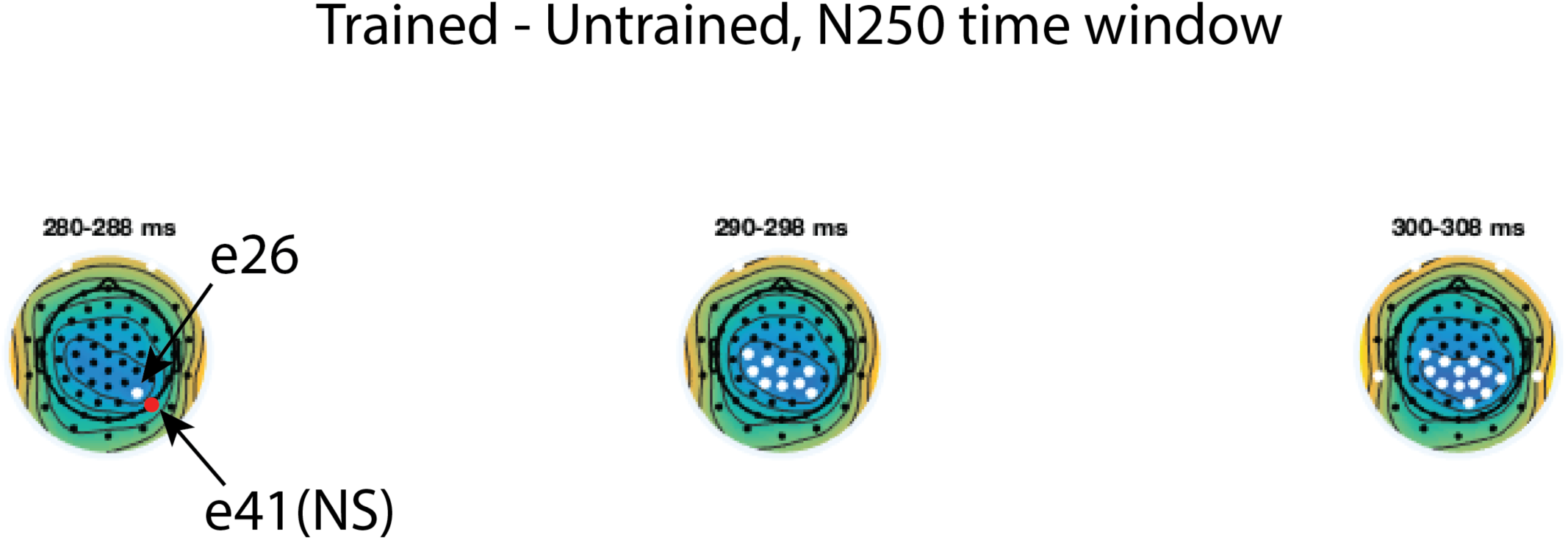
Closer view of the N250 time window from Figure S4.

**Figure S7.**
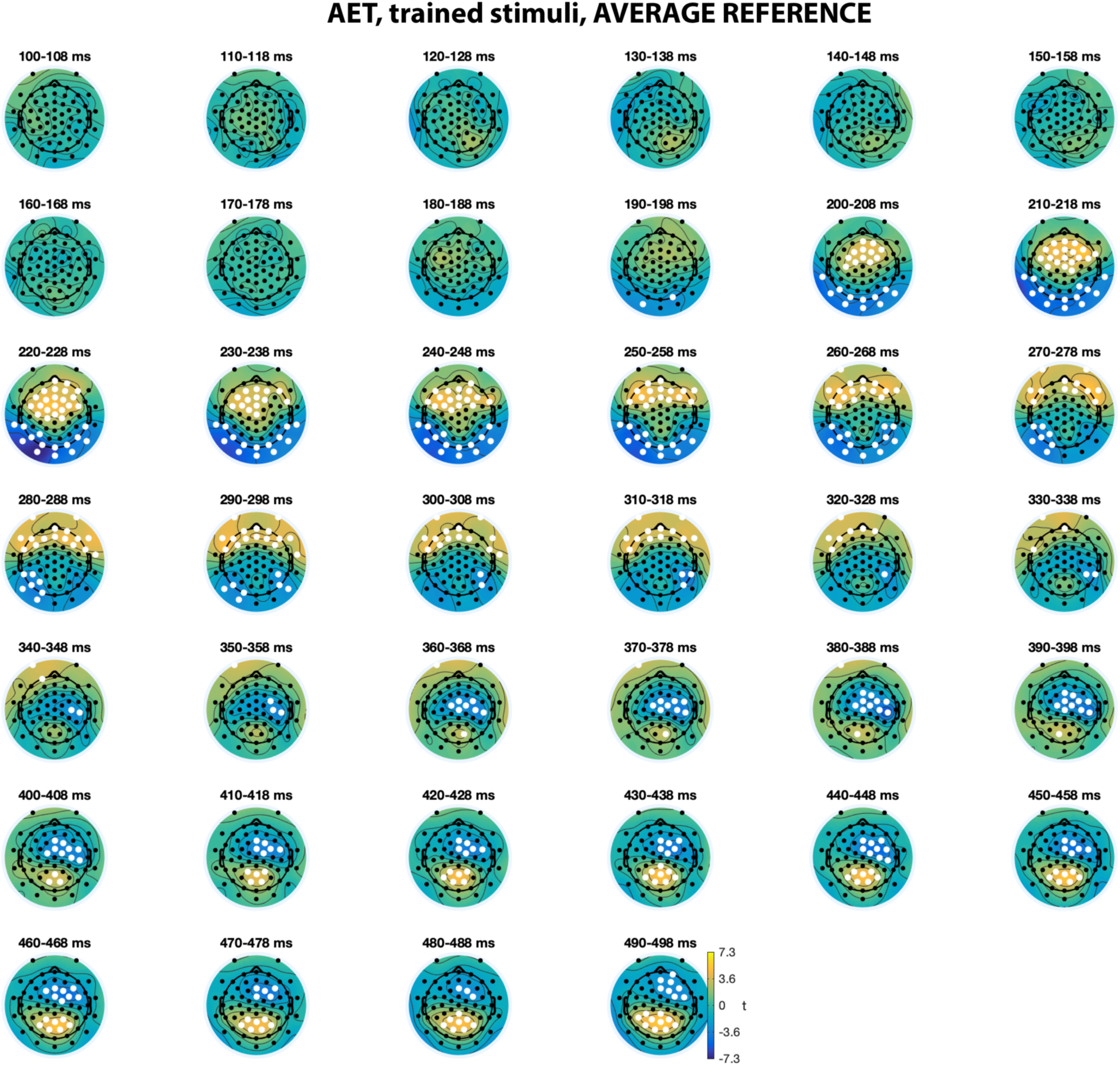
Mass univariate analysis of trained AET calculated from average reference data.

**Figure S8.**
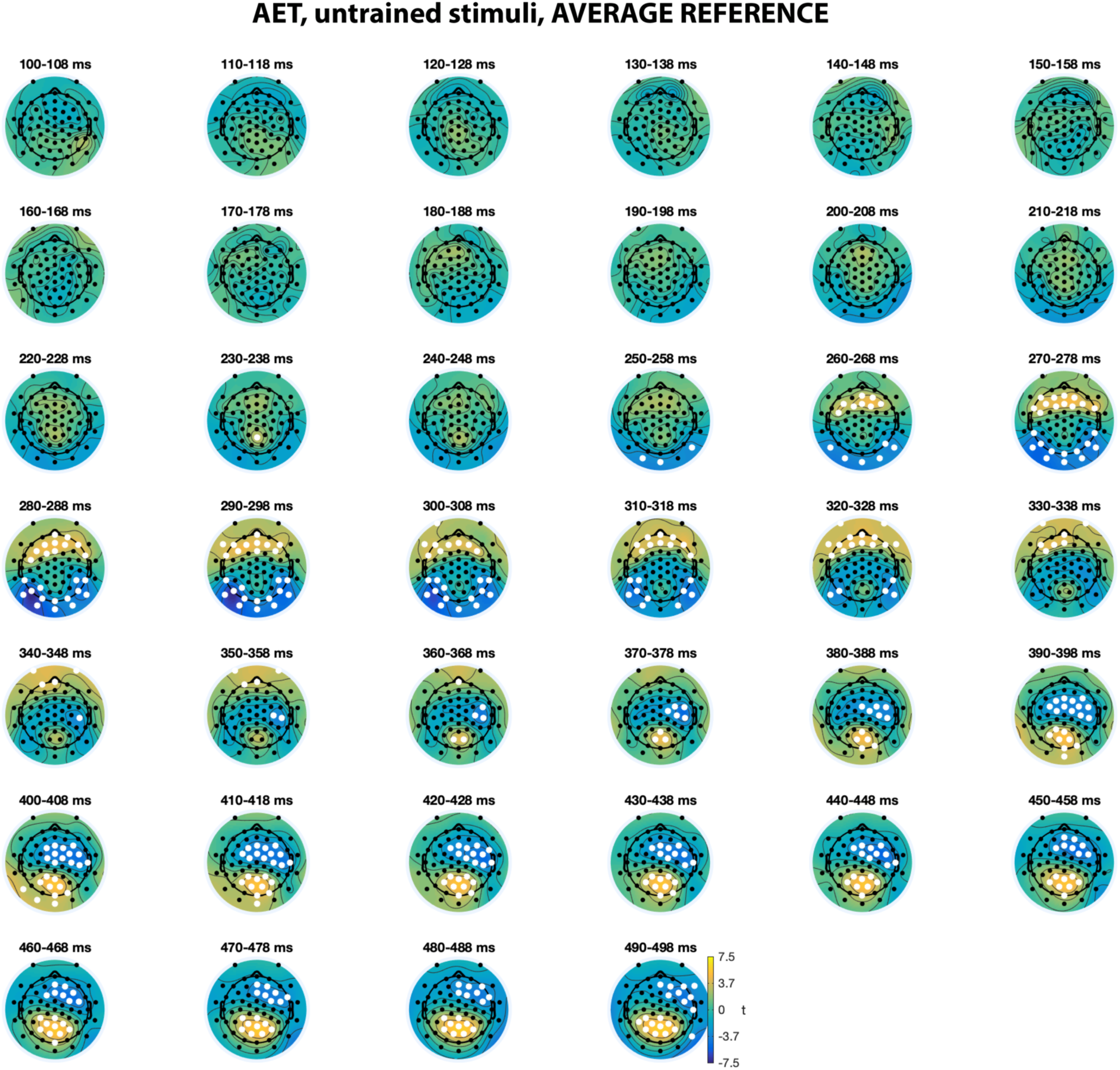
Mass univariate analysis of untrained AET calculated from average reference data.

**Figure S9.**
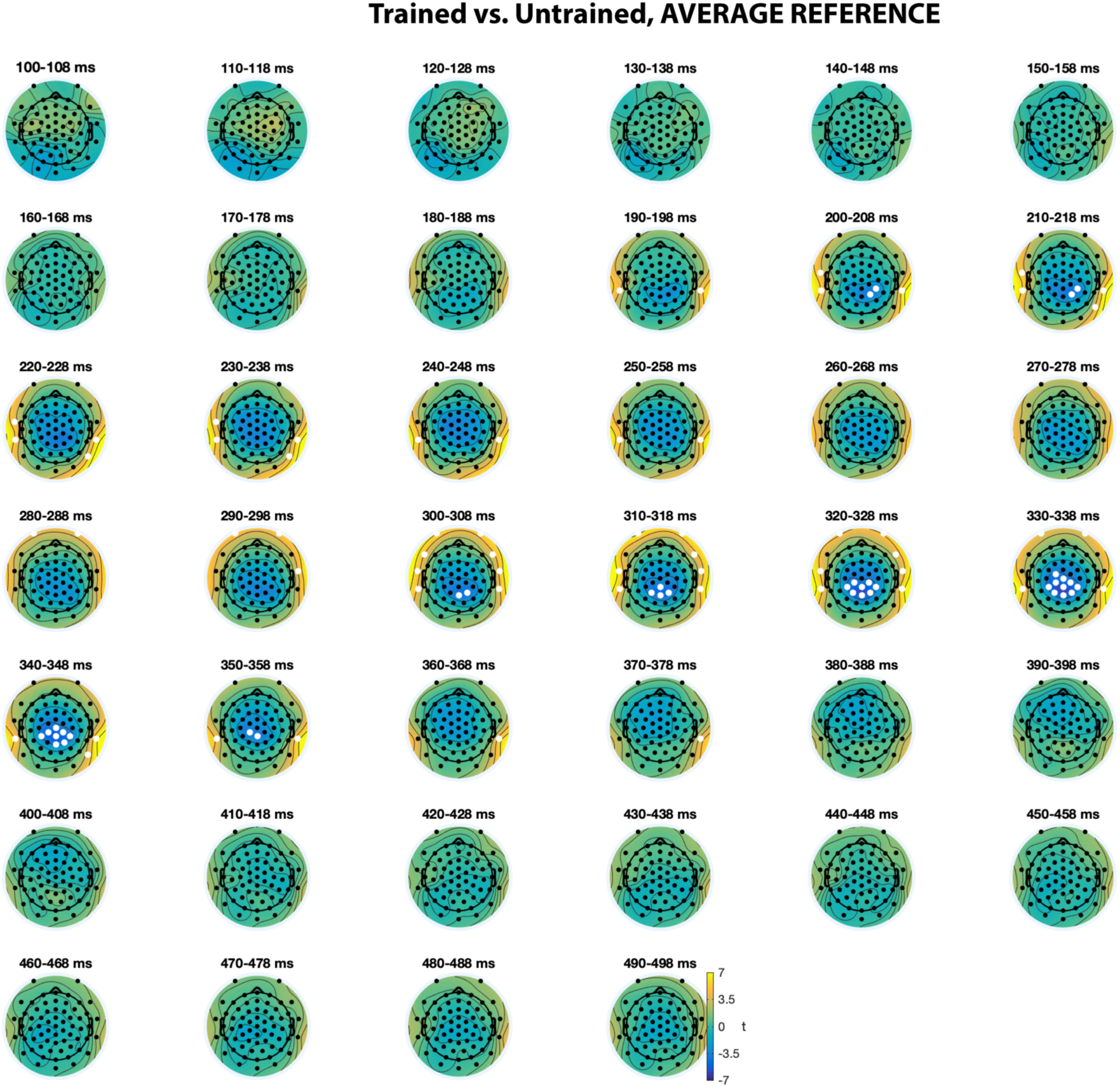
Mass univariate analysis of average reference trained vs. untrained difference wave. For the N200 epoch (190-260ms), the lateral positive difference is intact but the centro-parietal negative difference is greatly reduced relative to REST.

**Figure S10.**
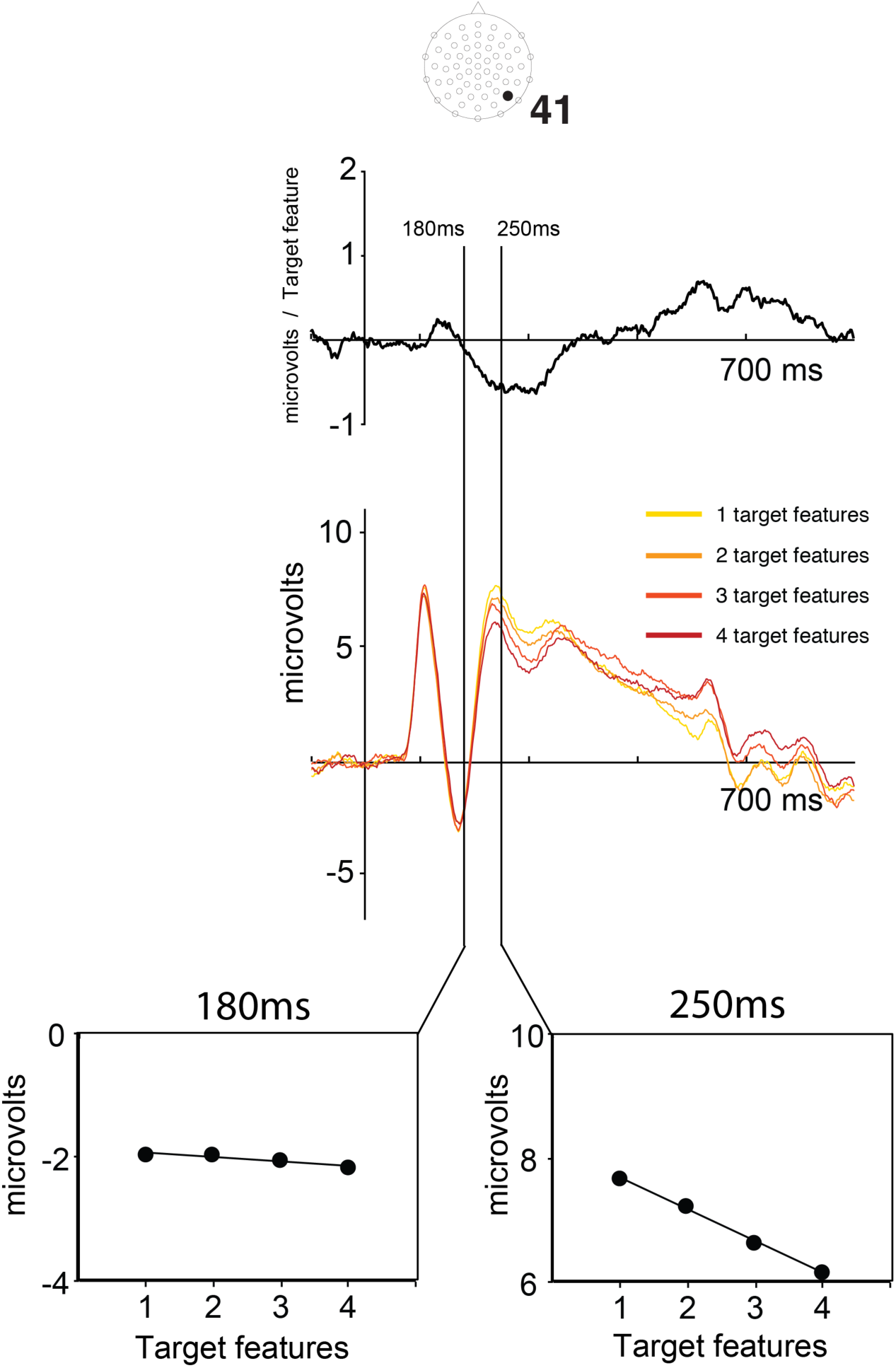
Construction of the attention effect timecourse for a sample electrode. Each timepoint at each electrode in each participant was measured for the 1, 2, 3, and 4 target feature conditions. The bottom panel shows these measurements for 180ms and 250ms. A line was then fit to the effect of target features (bottom panel). The slope of this line was plotted for each timepoint (top panel). Data in the bottom and middle panels are from the grand average. The AET in the top panel shows the grand average of individual participant AETs.

## Footnotes

1 We also tried fitting an exponential function to the SN, resulting in nearly identical timecourses and R^2^ values.

2 To evaluate the interaction, difference waves were calculated between the trained and untrained SN in each SOA condition and compared in the mass univariate analysis. No differences reached significance. See also, the supplementary latency analysis at electrodes 41 and 45, where no differences were found.

3 Recall that the mass univariate analysis did not reveal any differences between the trained and untrained AET, suggesting that there was no large interaction between the linear effect of training and the effect of target features. Comparison of training effects for the long and short SOAs in a supplementary mass univariate analysis [(Trained_LongSOA_ – Untrained_LongSOA_) – (Trained_ShortSOA_ – Untrained_ShortSOA_)] also revealed no evidence of an interaction between training and SOA.

## References

Allen, S. W., & Brooks, L. R. (1991). Specializing the operation of an explicit rule. Journal of Experimental Psychology: General, 120, 3–19.

Azizian, A., Freitas, A., Watson, T., & Squires, N. (2006). Electrophysiological correlates of categorization: P300 amplitude as index of target similarity. Biological Psychology, 71, 278–288.

Baldauf, D., & Desimone, R. (2014). Neural mechanisms of object-based attention. Science, 344, 424–427. doi: 10.1126/science.1247003

Bar, M. (2003). A cortical mechanism for triggering top-down facilitation in visual object recognition. Journal of Cognitive Neuroscience, 15, 600–609.

Barense, M. D., Ngo, J. K., Hung, L. H., & Peterson, M. A. (2012). Interactions of memory and perception in amnesia: The figure–ground perspective. Cerebral Cortex, 22, 2680–2691. doi: 10.1093/cercor/bhr347

Benjamini, Y., & Hochberg, Y. (1995). Controlling the false discovery rate: a practical and powerful approach to multiple testing. Journal of the royal statistical society. Series B (Methodological), 289–300.

Bentin, S., Mouchetant-Rostaing, Y., Giard, M.-H., Echallier, J.-F., & Pernier, J. (1999). ERP manifestations of processing printed words at different psycholinguistic levels: time course and scalp distribution. Journal of Cognitive Neuroscience, 11, 235–260.

Bilalić, M., Grottenthaler, T., Nägele, T., & Lindig, T. (2014). The faces in radiological images: fusiform face area supports radiological expertise. Cerebral Cortex, 26, 1004–1014.

Bilalić, M., Langner, R., Ulrich, R., & Grodd, W. (2011). Many faces of expertise: fusiform face area in chess experts and novices. The Journal of Neuroscience, 31, 10206–10214.

Blair, M. R., Watson, M. R., Walshe, R. C., & Maj, F. (2009). Extremely selective attention: eye-tracking studies of the dynamic allocation of attention to stimulus features in categorization. Journal of Experimental Psychology. Learning, Memory, and Cognition, 35, 1196–1206. doi: 2009-12193-008 [pii] 10.1037/a0016272

Carreiras, M., Duñabeitia, J. A., & Molinaro, N. (2009). Consonants and vowels contribute differently to visual word recognition: ERPs of relative position priming. Cerebral Cortex, 19, 2659–2670.

Chua, K.-W., Richler, J. J., & Gauthier, I. (2015). Holistic Processing From Learned Attention to Parts. Journal of Experimental Psychology: General, 144, 723–729. doi: 10.1037/xge0000063

Curran, T., & Cleary, A. M. (2003). Using ERPs to dissociate recollection from familiarity in picture recognition. Cognitive Brain Research, 15, 191–205.

Daffner, K. R., Zhuravleva, T. Y., Sun, X., Tarbi, E. C., Haring, A. E., Rentz, D. M., & Holcomb, P. J. (2012). Does modulation of selective attention to features reflect enhancement or suppression of neural activity? Biological Psychology, 89, 398–407.

Dehaene, S., & Cohen, L. (2011). The unique role of the visual word form area in reading. Trends in Cognitive Sciences, 15, 254–262.

Dieciuc, M., Roque, N. A., & Folstein, J. R. (2017). Changing similarity: Stable and flexible modulations of psychological dimensions. Brain Research, 1670, 208–219.

Dong, L., Li, F., Liu, Q., Wen, X., Lai, Y., Xu, P., & Yao, D. (2017). MATLAB toolboxes for reference electrode standardization technique (REST) of scalp EEG. Frontiers in neuroscience, 11, 601.

Du, Y., Hu, W., & Fang, Z. (2013). Electrophysiological correlates of morphological processing in Chinese compound word recognition. Frontiers in human neuroscience, 7, 601.

Du, Y., Zhang, Q., & Zhang, J. X. (2014). Does N200 reflect semantic processing?—An ERP study on Chinese visual word recognition. PLoS One, 9, e90794.

Folstein, J. R., Fuller, K., Howard, D., & DePatie, T. (2017). The effect of category learning on attentional modulation of visual cortex. Neuropsychologia, 104, 18–30.

Folstein, J. R., Monfared, S. S., & Maravel, T. (2017). The effect of category learning on visual attention and visual representation. Psychophysiology, 54, 1855–1871.

Folstein, J. R., Palmeri, T. J., Van Gulick, A., & Gauthier, I. (2015). Categoy learning stretches neural representations in visual cortex. Current Directions in Psychological Science, 24, 17–23.

Folstein, J. R., & Van Petten, C. (2004). Multidimensional rule, unidimensional rule, and similarity strategies in categorization: event-related potential correlates. Journal of Experimental Psychology: Learning, Memory, and Cognition, 30, 1026–1044.

Friston, K., & Kiebel, S. (2009). Predictive coding under the free-energy principle. Philosophical Transactions of the Royal Society of London B: Biological Sciences, 364, 1211–1221.

Ganis, G., & Kosslyn, S. M. (2007). Multiple mechanisms of top-down processing in vision Representation and brain (pp. 21–45): Springer.

Gauthier, I., Skudlarski, P., Gore, J. C., & Anderson, A. W. (2000). Expertise for cars and birds recruits brain areas involved in face recognition. Nature Neuroscience, 3, 191–197.

Gauthier, I., & Tarr, M. J. (1997). Becoming a “Greeble” expert: exploring mechanisms for face recognition. Vision Research, 37, 1673–1682.

Gauthier, I., Tarr, M. J., Anderson, A. W., Skudlarski, P., & Gore, J. C. (1999). Activation of the middle fusiform ‘face area’ increases with expertise in recognizing novel objects. Nature Neuroscience, 2, 568–573.

Gil-da-Costa, R., Stoner, G. R., Fung, R., & Albright, T. D. (2013). Nonhuman primate model of schizophrenia using a noninvasive EEG method. Proceedings of the National Academy of Sciences, 201312264.

Gledhill, D., Grimsen, C., Fahle, M., & Wegener, D. (2015). Human feature-based attention consists of two distinct spatiotemporal processes. Journal of Vision, 15, 8–8. doi: 10.1167/15.8.8

Gordon, I., & Tanaka, J. W. (2011). The role of name labels in the formation of face representations in event-related potentials. British Journal of Psychology, 102, 884–898. doi: 10.1111/j.2044-8295.2011.02064.x

Grainger, J., Kiyonaga, K., & Holcomb, P. J. (2006). The time course of orthographic and phonological code activation. Psychological Science, 17, 1021–1026.

Gregoriou, G. G., Gotts, S. J., Zhou, H., & Desimone, R. (2009). High-frequency, long-range coupling between prefrontal and visual cortex during attention. Science, 324, 1207–1210. doi: 10.1126/science.1171402

Groppe, D. M., Urbach, T. P., & Kutas, M. (2011). Mass univariate analysis of event - related brain potentials/fields I: A critical tutorial review. Psychophysiology, 48, 1711–1725. doi: 10.1111/j.1469-8986.2011.01273.x

Harel, A., Gilaie-Dotan, S., Malach, R., & Bentin, S. (2010). Top-down engagement modulates the neural expressions of visual expertise. Cerebral Cortex, 20, 2304–2318. doi: 10.1093/cercor/bhp316

Herzmann, G. (2016). Increased N250 amplitudes for other-race faces reflect more effortful processing at the individual level. International Journal of Psychophysiology, 105, 57–65.

Hillyard, S. A., & Anllo-Vento, L. (1998). Event-related brain potentials in the study of visual selective attention. Proceedings of the National Academy of Sciences, 95, 781–787.

Holcomb, P. J., & Grainger, J. (2007). Exploring the temporal dynamics of visual word recognition in the masked repetition priming paradigm using event-related potentials. Brain Research, 1180, 39–58.

Jolicoeur, P., Gluck, M. A., & Kosslyn, S. M. (1984). Pictures and names: Making the connection. Cognitive Psychology, 16, 243–275.

Jones, T., Hadley, H., Cataldo, A. M., Arnold, E., Curran, T., Tanaka, J. W., & Scott, L. S. (2018). Neural and behavioral effects of subordinate-level training of novel objects across manipulations of color and spatial frequency. European Journal of Neuroscience.

Kiesel, A., Miller, J., Jolicœur, P., & Brisson, B. (2008). Measurement of ERP latency differences: A comparison of single - participant and jackknife - based scoring methods. Psychophysiology, 45, 250–274.

Lamme, V. A., & Roelfsema, P. R. (2000). The distinct modes of vision offered by feedforward and recurrent processing. Trends in Neurosciences, 23, 571–579.

Li, S., Ostwald, D., Giese, M., & Kourtzi, Z. (2007). Flexible coding for categorical decisions in the human brain. Journal of Neuroscience, 27, 12321–12330.

Liu, Q., Balsters, J. H., Baechinger, M., van der Groen, O., Wenderoth, N., & Mantini, D. (2015). Estimating a neutral reference for electroencephalographic recordings: the importance of using a high-density montage and a realistic head model. Journal of neural engineering, 12, 056012.

Maurer, U., Blau, V. C., Yoncheva, Y. N., & McCandliss, B. D. (2010). Development of visual expertise for reading: rapid emergence of visual familiarity for an artificial script. Developmental Neuropsychology, 35, 404–422.

McCandliss, B. D., Posner, M. I., & Givon, T. (1997). Brain plasticity in learning visual words. Cognitive Psychology, 33, 88–110.

McGugin, R. W., Gatenby, J. C., Gore, J. C., & Gauthier, I. (2012). High-resolution imaging of expertise reveals reliable object selectivity in the fusiform face area related to perceptual performance. Proceedings of the National Academy of Sciences, 109, 17063–17068.

McKone, E., Kanwisher, N., & Duchaine, B. C. (2007). Can generic expertise explain special processing for faces? Trends in Cognitive Sciences, 11, 8–15.

Medin, D. L., & Schaffer, M. M. (1978). Context theory of classification learning. Psychological Review, 85, 207–238.

Mei, L., Xue, G., Lu, Z.-L., He, Q., Zhang, M., Xue, F., … Dong, Q. (2013). Orthographic transparency modulates the functional asymmetry in the fusiform cortex: An artificial language training study. Brain and Language, 125, 165–172.

Moore, M. W., Durisko, C., Perfetti, C. A., & Fiez, J. A. (2014). Learning to read an alphabet of human faces produces left-lateralized training effects in the fusiform gyrus. Journal of Cognitive Neuroscience, 26, 896–913.

Mulert, C., Jäger, L., Schmitt, R., Bussfeld, P., Pogarell, O., Möller, H.-J., … Hegerl, U. (2004). Integration of fMRI and simultaneous EEG: towards a comprehensive understanding of localization and time-course of brain activity in target detection. Neuroimage, 22, 83–94.

Pascual-Marqui, R. D. (2002). Standardized low-resolution brain electromagnetic tomography (sLORETA): technical details. Methods and Findings in Experimental and Clinical Pharmacology, 24, 5–12.

Pascual-Marqui, R. D. (2009). Theory of the EEG inverse problem. Quantitative EEG analysis: methods and clinical applications, 121–140.

Pierce, L. J., Scott, L. S., Boddington, S., Droucker, D., Curran, T., & Tanaka, J. W. (2011). The N250 brain potential to personally familiar and newly learned faces and objects. Frontiers in human neuroscience, 5. doi: 10.3389/fnhum.2011.00111

Price, C. J., & Devlin, J. T. (2011). The interactive account of ventral occipitotemporal contributions to reading. Trends in Cognitive Sciences, 15, 246–253.

Ruz, M., & Nobre, A. C. (2008). Attention modulates initial stages of visual word processing. Journal of Cognitive Neuroscience, 20, 1727–1736.

Schendan, H. E., & Ganis, G. (2015). Top-down modulation of visual processing and knowledge after 250 ms supports object constancy of category decisions. Frontiers in Psychology, 6. doi: 10.3389/fpsyg.2015.01289

Schendan, H. E., & Kutas, M. (2003). Time course of processes and representations supporting visual object identification and memory. Journal of Cognitive Neuroscience, 15, 111–135.

Schendan, H. E., & Lucia, L. C. (2010). Object-sensitive activity reflects earlier perceptual and later cognitive processing of visual objects between 95 and 500 ms. Brain Research, 1329, 124–141.

Schendan, H. E., & Maher, S. M. (2009). Object knowledge during entry-level categorization is activated and modified by implicit memory after 200 ms. Neuroimage, 44, 1423–1438.

Schweinberger, S. R., & Neumann, M. F. (2016). Repetition effects in human ERPs to faces. Cortex, 80, 141–153.

Scott, L., Tanaka, J. W., Sheinberg, D. L., & Curran, T. (2006). A reevaluation of the electrophysiological correlates of expert object processing. Cognitive Neuroscience, Journal of, 18, 1453–1465. doi: 10.1162/jocn.2006.18.9.1453

Scott, L., Tanaka, J. W., Sheinberg, D. L., & Curran, T. (2008). The role of category learning in the acquisition and retention of perceptual expertise: A behavioral and neurophysiological study. Brain Research, 1210, 204–215. doi: 10.1016/j.brainres.2008.02.054

Smid, H. G., Jakob, A., & Heinze, H.-J. (1999). An event-related brain potential study of visual selective attention to conjunctions of color and shape. Psychophysiology, 36, 264–279. doi: 10.1017/S0048577299971135

Song, Y., Hu, S., Li, X., Li, W., & Liu, J. (2010). The role of top-down task context in learning to perceive objects. Journal of Neuroscience, 30, 9869–9876. doi: 10.1523/JNEUROSCI.0140-10.2010

Takahashi, H., Rissling, A. J., Pascual-Marqui, R., Kirihara, K., Pela, M., Sprock, J., … Light, G. A. (2013). Neural substrates of normal and impaired preattentive sensory discrimination in large cohorts of nonpsychiatric subjects and schizophrenia patients as indexed by MMN and P3a change detection responses. Neuroimage, 66, 594–603.

Tanaka, J. W., & Taylor, M. (1991). Object categories and expertise: Is the basic level in the eye of the beholder? Cognitive Psychology, 23, 457–482.

Ulrich, R., & Miller, J. (2001). Using the jackknife-based scoring method for measuring LRP onset effects in factorial designs. Psychophysiology, 38, 816–827.

Wong, A. C., Palmeri, T. J., & Gauthier, I. (2009). Conditions for facelike expertise with objects: becoming a Ziggerin expert--but which type? Psychol Sci, 20, 1108–1117. doi: PSCI2430 [pii] 10.1111/j.1467-9280.2009.02430.x

Xue, G., Chen, C., Jin, Z., & Dong, Q. (2006). Language experience shapes fusiform activation when processing a logographic artificial language: an fMRI training study. Neuroimage, 31, 1315–1326.

Yao, D. (2001). A method to standardize a reference of scalp EEG recordings to a point at infinity. Physiological Measurement, 22, 693.

Yoncheva, Y. N., Blau, V. C., Maurer, U., & McCandliss, B. D. (2010). Attentional focus during learning impacts N170 ERP responses to an artificial script. Developmental Neuropsychology, 35, 423–445.

Yoncheva, Y. N., Wise, J., & McCandliss, B. (2015). Hemispheric specialization for visual words is shaped by attention to sublexical units during initial learning. Brain and Language, 145, 23–33.

Yovel, G., Yovel, I., & Levy, J. (2001). Hemispheric asymmetries for global and local visual perception: effects of stimulus and task factors. Journal of Experimental Psychology: Human Perception and Performance, 27, 1369.

Zhang, J. X., Fang, Z., Du, Y., Kong, L., Zhang, Q., & Xing, Q. (2012). Centro-parietal N200: an event-related potential component specific to Chinese visual word recognition. Chinese Science Bulletin, 57, 1516–1532.

Zhou, A., Yin, Y., Zhang, J., & Zhang, R. (2016). Does font type influence the N200 enhancement effect in Chinese word recognition? Journal of Neurolinguistics, 39, 57–68.

